# Classical guinea pig coat phenotypes are associated with variants in *FGF5* and *CELSR1*

**DOI:** 10.64898/2026.11.02.763795

**Authors:** Madalene Halley, Craig J. Bierle, PingHsun Hsieh, Arslan Zaidi, Frank W. Albert, Maureen Cetera

## Abstract

Classical Mendelian coat phenotypes in the domestic guinea pig, *Cavia porcellus*, were central to early studies of mammalian genetics, yet their molecular bases remain unresolved. Long fur is characterized by elongated hair, and rough coat by whorled patterns of fur orientation. Here, we performed genome-wide association mapping using exome and whole-genome sequencing data to identify the genetic basis of both phenotypes. Mapping of long fur identified a locus containing *FGF5* and resolved the association to a premature stop codon, independently confirming a previously proposed candidate variant. Mapping of rough coat identified a locus containing *CELSR1*, an atypical cadherin and core component of the planar cell polarity (PCP) pathway, resolving the association to an in-frame proline insertion in its extracellular domain. The variant is associated with a dominant rough-coat phenotype, unlike previously characterized recessive mouse *Celsr1* alleles, which produce a similar hair-patterning phenotype only when homozygous. The mouse alleles are also associated with severe developmental abnormalities, including open neural tube, which are not observed in rough-coat guinea pigs. This dominant, tissue-specific phenotype suggests that the insertion alters CELSR1 function in a previously undescribed manner. Consistent with this possibility, the rough-coat variant localized normally to cell-cell junctions in cultured mouse keratinocytes and retained its ability to recruit Fzd6 and Vangl2, demonstrating preserved molecular interactions involved in PCP junction assembly. These findings resolve the molecular basis of two classical guinea pig coat phenotypes and identify a naturally occurring, dominant *CELSR1* allele associated with a mode of PCP disruption distinct from previously characterized alleles.

## INTRODUCTION

The domestic guinea pig (*Cavia porcellus*) has a long history as a model for mammalian genetics (Pritt, 2012). This work benefited from numerous established guinea pig varieties with distinctive, heritable coat phenotypes that were maintained by animal fanciers before the adoption of guinea pigs as an experimental model (Cumberland, 1886). Among the most distinctive breeds are Abyssinian, characterized by a rough coat, and Peruvian, which combines the rough coat with long fur; both breeds were documented as early as 1886 (Castle & Wright, 1916; Cumberland, 1886). By the early twentieth century, rough coat and long fur were recognized as Mendelian traits, with rough coat inherited dominantly and long fur inherited recessively (Castle, 1903, 1905). Sewall Wright subsequently conducted extensive breeding studies of guinea pig coat phenotypes, documenting inheritance patterns and genetic interactions that contribute to variation in coat color and coat texture (Castle & Wright, 1916; Wright, 1927, 1935, 1949, 1984). These studies established the guinea pig as a powerful system for investigating how phenotypic variation can arise from individual loci and interactions among genetic factors. However, despite more than a century of phenotypic and genetic characterization, the molecular basis of these classical coat traits has remained unresolved.

To this day, breeders have continued to maintain classical coat varieties and the distinctive phenotypes on which they were originally defined. Long fur results in substantially elongated hair, while rough coat is characterized by whorled rosette patterns of fur orientation distributed across the body, in contrast to the uniformly oriented fur of smooth coat, wild-type guinea pigs (Figure 1) (Castle & Wright, 1916; Wright, 1984). A candidate causal variant in *FGF5* was previously proposed for long fur based on targeted sequencing of a strong biological candidate, consistent with the conserved role of *FGF5* as a negative regulator of hair growth across mammals (Carrion et al., 2024; Yu et al., 2018). However, this candidate has not been confirmed using an unbiased genome-wide mapping approach. In contrast, the genetic basis of rough coat has never been identified.

**Figure 1.**
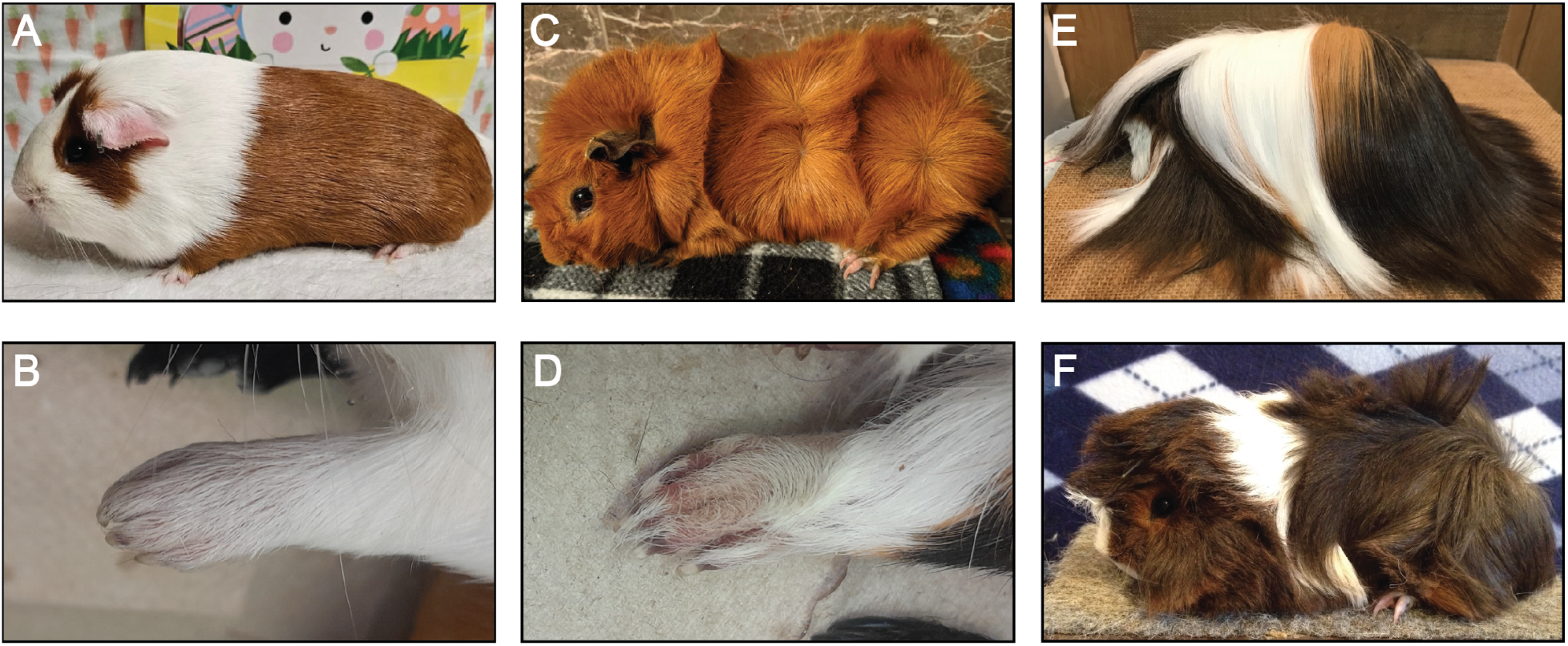
Coat characteristics of American, Abyssinian, and Peruvian guinea pig breeds. (A-B) American guinea pigs have a smooth coat with short fur and uniformly oriented fur on the paws (B). (C-D) Abyssinian guinea pigs have a rough coat with short fur and misoriented fur on the paws (D). (E-F) Peruvian guinea pigs have long fur and a rough coat, which is apparent in juveniles before the fur reaches full length (F).

Unlike long fur, the presentation of rough coat can vary considerably in the completeness and distribution of rosette patterns. This variability was itself a major focus of Wright’s genetic studies, which established a primary locus for rough coat and the existence of additional loci that modify the extent and distribution of rosettes (Wright, 1935, 1949, 1950, 1984). Despite this complexity, the presence or absence of rough coat reflects a binary state dependent on a single locus.

The whorled hair orientations in rough-coat guinea pigs are reminiscent of hair-patterning phenotypes in mice caused by disrupting the planar cell polarity (PCP) pathway, a highly conserved pathway that establishes tissue-wide polarity to direct cellular behaviors during development (Basta et al., 2022; Butler & Wallingford, 2017; Cetera et al., 2017; Guo et al., 2004; Ravni et al., 2009). *Celsr1* encodes an atypical cadherin and core component of the PCP machinery. Celsr1 proteins on neighboring cells bind one another across the cell-cell junction. The other core transmembrane PCP proteins, Fzd6 and Vangl2, associate with Celsr1 on opposing sides of the junction, establishing an asymmetric PCP complex between neighboring cells (Butler & Wallingford, 2017; Devenport, 2014; Harrison et al., 2020). Coordination of these complexes across the tissue establishes a common axis of cellular orientation (Butler & Wallingford, 2017). In the skin, the tissue-wide axis governs the coordinated orientation of hair follicles (Devenport & Fuchs, 2008).

Unlike rough coat, which is dominant and fully penetrant (Wright, 1949), comparable hair-patterning defects in mice occur only in animals homozygous for *Celsr1* loss-of-function alleles, while heterozygotes show no phenotype (Basta et al., 2022; Ravni et al., 2009). *Celsr1* null alleles produce variable whorled hair patterns in homozygous mice, but these animals can also suffer from lethal open neural tube defects, and those that survive are smaller and less healthy than littermate controls (Basta et al., 2022; Ravni et al., 2009). Other core PCP alleles, including *Vangl2^ΔTm^*, *Vangl2^Lp^*, and *Celsr1^Crsh^*, cause near-fully penetrant neural tube defects and lethality in homozygotes, preventing the assessment of postnatal coat phenotypes (Curtin et al., 2003; Kibar et al., 2001; Murdoch et al., 2001; Yin et al., 2012). *Vangl2^ΔTm^* is a loss-of-function allele, while *Vangl2^Lp^*and *Celsr1^Crsh^* are missense mutations that are considered dominant-negative in some tissues, though heterozygotes do not show altered hair follicle orientation (Devenport & Fuchs, 2008; Ravni et al., 2009). These developmental defects have motivated conditional knockout approaches to assess postnatal hair follicle orientation. Combined loss of both *Vangl* paralogs, *Vangl1* and *Vangl2*, in the skin eliminates epidermal PCP activity and produces a whorled fur phenotype that closely resembles rough coat, without the neural tube defects or other systemic abnormalities associated with PCP disruption (Cetera et al., 2017).

The guinea pig rough coat therefore presents a striking genetic paradox: its phenotype resembles a severe disruption of PCP activity in the skin, yet the trait is dominant and occurs in otherwise healthy animals. These observations raised the possibility that naturally occurring variation could disrupt PCP-dependent hair patterning through a mechanism that is distinct from canonical loss of function.

Here, we investigated the genetic basis of rough coat and long fur in the domestic guinea pig using genome-wide association mapping. Using a combination of exome capture and whole-genome sequencing, mapping of long fur recovered a previously described premature stop codon in *FGF5*, providing independent, unbiased confirmation of the previously proposed candidate (Yu et al., 2018). For rough coat, we identified an in-frame proline insertion in the extracellular domain of CELSR1 as the likely causal variant. This variant is associated with a dominant PCP-related hair-patterning phenotype that is distinct from those produced by previously characterized PCP disruptions in mice (Basta et al., 2022; Curtin et al., 2003; Kibar et al., 2001; Murdoch et al., 2001; Ravni et al., 2009; Yin et al., 2012). In culture, the proline insertion does not alter protein localization or recruitment of other core PCP proteins, suggesting the variant is not a simple loss-of-function. Together, these findings identify the likely molecular basis of two classical guinea pig coat traits and identify a naturally occurring, dominant *CELSR1* variant that may represent a distinct class of PCP alleles.

## RESULTS

### Rough coat and long fur are classic guinea pig coat phenotypes

Rough coat exhibits substantial variation in the extent and distribution of rosette patterns. At one extreme, full rough animals have whorled rosette patterns of hair orientation distributed across the body, including the paws, whereas at the other extreme, altered hair direction is restricted to the paws (Wright, 1949). The full rough phenotype is characteristic of the Abyssinian breed, in contrast to the uniformly oriented coat of smooth animals (Figure 1A-D). The Peruvian guinea pig breed combines the rough-coat phenotype with long fur. The long fur of these animals obscures the underlying rosette pattern (Figure 1E), but rosettes are apparent in juveniles before the fur reaches its full length (Figure 1F). Although the number and position of rosettes vary among rough-coat animals, rosettes were apparent in all animals classified as rough in this study. For genetic mapping, we therefore scored rough coat as a binary phenotype based on the presence or absence of rosettes, and long fur as a binary phenotype based on hair length.

### Genetic mapping identifies a single association locus containing *FGF5* for recessive long fur

We sequenced 54 guinea pigs using the following platforms: human exome (Illumina), n = 32; whole-genome (Illumina), n = 2; whole-genome (Ultima), n = 20. Because no guinea pig-specific exome capture kit is commercially available, we performed exome capture using a human exome capture kit and sequenced the resulting libraries. An average of 22 million total reads per sample were generated, of which 93% mapped confidently (map quality ≥ 30) to the guinea pig reference genome mCavPor4.1 (NCBI RefSeq GCF_034190915.1). After filtering for reads with a map quality ≥ 30, exome-captured samples had an average depth of 33.2x over the guinea pig coding sequence. For whole-genome sequencing samples, Illumina samples had an average sequencing depth of 34.2x, and Ultima samples had an average depth of 37.1x.

Using these data, genome-wide association mapping was performed using a linear mixed model implemented in GEMMA (X. Zhou & Stephens, 2012), incorporating sequenced animals and the phenotypes recorded in Table 1: 14 long-fur, 35 short-fur, and 5 with an unknown phenotype because the animals were too young to assess final coat length (Figure S1). Most samples were obtained from a single breeder, with additional samples from local shelters and pet owners. Relatedness among animals was therefore expected due to shared breeding histories, and a centered kinship matrix was incorporated into the association model to account for population structure. GEMMA excludes variants missing genotype calls in more than 5% of samples; because most samples were exome-sequenced, this restricted the analysis primarily to coding variants, yielding 386,066 variants for association testing.

**Table 1.** Sequencing, phenotype, and genotype information for guinea pig samples. For the long-fur phenotype, samples are labeled long or short based on coat length. Samples labeled with unkn (unknown abbreviation) long-fur phenotype are pups without fully grown coats. For the rough-coat phenotype, samples are labeled smooth or rough to represent fur whorl presence or absence. Samples labeled with unkn were not well preserved and the presence or absence of whorls was not able to be determined. For genotype information, 0 indicates the reference allele while 1 indicates the alternative allele.

| **Sample** | **Sequencing** | **Rough Phenotype** | **XM_013158622.2:c.3842_3843insGCC** | **Fur Length Phenotype** | **XM_003477669.5:c.403C>T** | **Notes** |
|---|---|---|---|---|---|---|
| **gp1** | WGSIllumina | Rough | 1/1 | short | 0/0 | Sequencing labeled Aby or AbyTissue |
| **gp2** | WGSIllumina | unkn | 1/1 | long | 1/1 | Sequencing labeled Peru or PeruTissue |
| **gp3** | WGSUltima | Rough | 0/1 | unkn | 0/1 | pup |
| **gp4** | WGSUltima,Sanger | Rough | 0/1 | unkn | 0/1 | pup |
| **gp5** | ExomeIllumina | Rough | 0/1 | short | 0/0 |  |
| **gp6** | ExomeIllumina | Rough | 1/1 | short | 0/0 |  |
| **gp7** | ExomeIllumina | Rough | 1/1 | short | 0/0 |  |
| **gp8** | ExomeIllumina | Rough | 1/1 | short | 0/0 |  |
| **gp9** | ExomeIllumina | Rough | 0/1 | short | 0/0 |  |
| **gp10** | ExomeIllumina | Rough | 1/1 | short | 0/0 |  |
| **gp11** | ExomeIllumina | Rough | 1/1 | short | 0/0 |  |
| **gp12** | ExomeIllumina | Rough | 1/1 | short | 0/0 |  |
| **gp13** | ExomeIllumina | Rough | 1/1 | short | 0/0 |  |
| **gp14** | ExomeIllumina | Smooth | 0/0 | short | 0/0 |  |
| **gp15** | ExomeIllumina,Sanger | Smooth | 0/0 | short | 0/0 |  |
| **gp16** | ExomeIllumina | Smooth | 0/0 | short | 0/0 |  |
| **gp17** | ExomeIllumina | Smooth | 0/0 | short | 0/0 |  |
| **gp18** | WGSUltima | Smooth | 0/0 | long | 1/1 |  |
| **gp19** | WGSUltima | Rough | 0/1 | unkn | 1/1 | pup |
| **gp24** | ExomeIllumina | Smooth | 0/0 | long | 1/1 |  |
| **gp25** | ExomeIllumina,Sanger | unkn | 0/1 | long | 1/1 |  |
| **gp26** | ExomeIllumina | Rough | 1/1 | short | 0/0 |  |
| **gp27** | ExomeIllumina,Sanger | Rough | 0/1 | short | 0/0 |  |
| **gp28** | ExomeIllumina | Rough | 1/1 | short | 0/0 |  |
| **gp29** | ExomeIllumina,Sanger | Smooth | 0/0 | short | 0/1 |  |
| **gp43** | WGSUltima | Rough | 0/1 | short | 0/0 |  |
| **gp44** | ExomeIllumina | Rough | 0/1 | short | 0/1 |  |
| **gp45** | ExomeIllumina | Smooth | 0/0 | short | 0/1 |  |
| **gp46** | ExomeIllumina | Smooth | 0/0 | unkn | 0/0 | pup |
| **gp47** | ExomeIllumina | Rough | 0/1 | short | 0/1 |  |
| **gp48** | ExomeIllumina,Sanger | Smooth | 0/0 | short | 0/1 |  |
| **gp49** | ExomeIllumina | Smooth | 0/0 | short | 0/0 |  |
| **gp50** | ExomeIllumina | unkn | 0/0 | long | 1/1 |  |
| **gp51** | ExomeIllumina,Sanger | Rough | 0/1 | short | 0/0 |  |
| **gp52** | ExomeIllumina | Rough | 0/1 | short | 0/0 |  |
| **gp53** | ExomeIllumina | Smooth | 0/0 | short | 0/0 |  |
| **gp55** | WGSUltima,Sanger | Smooth | 0/0 | short | 0/0 |  |
| **gp56** | ExomeIllumina | Rough | 0/1 | long | 1/1 |  |
| **gp57** | ExomeIllumina | Smooth | 0/0 | short | 0/1 |  |
| **gp58** | WGSUltima | Smooth | 0/0 | long | 1/1 |  |
| **gp59** | WGSUltima | Smooth | 0/0 | long | 1/1 |  |
| **gp60** | ExomeIllumina | Smooth | 0/0 | short | 0/1 |  |
| **gp64** | WGSUltima,Sanger | Rough | 0/1 | unkn | 1/1 | pup |
| **gp65** | WGSUltima,Sanger | Smooth | 0/0 | long | 1/1 |  |
| **gp66** | Sanger | Rough | 1/1 | short | 0/0 |  |
| **gp68** | WGSUltima,Sanger | Rough | 0/1 | short | 0/0 |  |
| **gp69** | Sanger | Rough | 0/1 | - | - |  |
| **gp70** | Sanger | Smooth | 0/0 | - | - |  |
| **gp71** | Sanger | Rough | 1/1 | short | 0/0 |  |
| **gp72** | Sanger | Rough | 1/1 | short | 0/0 |  |
| **gp73** | Sanger | Rough | 1/1 | short | 0/0 |  |
| **gp74** | Sanger | Smooth | 0/0 | short | 0/0 |  |
| **gp75** | WGSUltima,Sanger | Rough | 0/1 | short | 0/1 |  |
| **gp76** | Sanger | Smooth | 0/0 | short | 0/0 |  |
| **gp77** | WGSUltima,Sanger | unkn | 0/0 | long | 1/1 |  |
| **gp78** | Sanger | Smooth | 0/0 | short | 0/0 |  |
| **gp79** | Sanger | Smooth | 0/0 | short | 0/0 |  |
| **gp80** | Sanger | Smooth | 0/0 | short | 0/0 |  |
| **gp81** | Sanger | Smooth | 0/0 | short | 0/0 |  |
| **gp82** | Sanger | Smooth | 0/0 | short | 0/0 |  |
| **gp85** | Sanger | Rough | 1/1 | - | - |  |
| **gp86** | WGSUltima | Smooth | 0/0 | long | 1/1 |  |
| **gp88** | Sanger | - | - | long | 1/1 |  |
| **gp89** | Sanger | - | - | long | 1/1 |  |
| **gp94** | WGSUltima | Smooth | 0/0 | long | 1/1 |  |
| **gp96** | Sanger | - | - | long | 1/1 |  |
| **gp104** | WGSUltima | Rough | 0/1 | short | 0/0 |  |
| **gp108** | Sanger | - | - | long | 1/1 |  |
| **gp109** | Sanger | - | - | long | 1/1 |  |
| **gp111** | WGSUltima | Rough | 0/1 | short | 0/0 |  |
| **gp113** | WGSUltima | Smooth | 0/0 | long | 1/1 |  |
| **gp116** | WGSUltima | Rough | 0/1 | short | 0/0 |  |
| **gp121** | WGSUltima | Smooth | 0/0 | long | 1/1 |  |
| **gp130** | Sanger | - | - | long | 1/1 |  |
| **gp135** | Sanger | - | - | long | 1/1 |  |
| **gp139** | Sanger | - | - | long | 1/1 |  |
| **gp140** | Sanger | - | - | long | 1/1 |  |
| **gp141** | Sanger | - | - | long | 1/1 |  |
| **gpC** | Sanger | Smooth | 0/0 | - | - | Short hair S2 or S13 lab strain |

Twelve variants across four scaffolds showed associations with long fur that exceeded the Bonferroni-corrected significance threshold (-log10(p) = 6.89; Figure 2A, Table 2), with no evidence of genomic inflation in the corresponding Q-Q plot (λ = 0.943; Figure 2B) and effect sizes (β) indicating a 0.30-0.72 increase in the predicted probability of the long-fur phenotype per additional effect allele (Figure 2C). Linkage disequilibrium between significant variants on the same scaffold was high on NW_026947511.1 (r = 0.88–0.90), a relatively compact ∼430 kb region. On NW_026947487.1, which spans a much larger ∼14.1 Mb region, linkage disequilibrium was strongest among more closely positioned variants and weaker between more distant variant pairs (r = 0.45–0.99; Figure 2D), consistent with expected LD decay across a wider physical distance.

**Figure 2.**
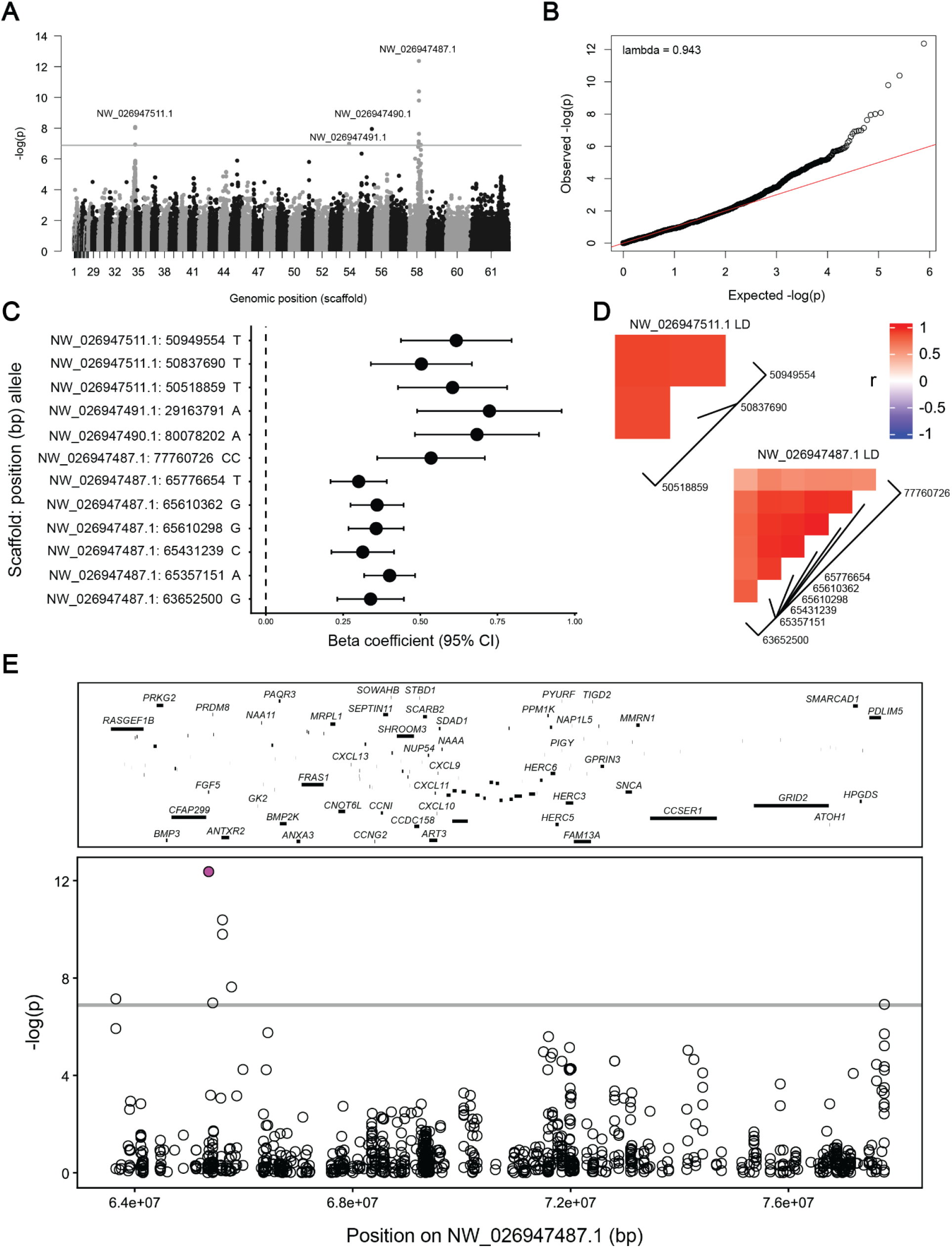
Genome-wide association mapping identifies a locus containing *FGF5* associated with long fur. (A) Manhattan plot of genome-wide association results from GEMMA. Each point represents a variant, plotted by scaffold (numbered in descending order of size; full names are provided in Table 2) and association significance (−log₁₀(p)). The grey line indicates the Bonferroni-corrected significance threshold (−log₁₀(p) = 6.89). Significant variants occur on scaffolds 34 (NW_026947511.1), 54 (NW_026947491.1), 55 (NW_026947490.1), and 58 (NW_026947487.1). (B) Quantile–quantile (Q–Q) plot of observed versus expected −log₁₀(p) values (λ = 0.943). (C) β coefficients for significant variants, with scaffold, position (bp), and allele indicated on the y-axis and 95% confidence intervals shown. (D) Linkage disequilibrium (LD) among significant variants on scaffolds NW_026947511.1 and NW_026947487.1. Colors indicate pairwise correlation coefficients (*r*), from −1 (blue) through 0 (white) to 1 (red); variant positions (bp) are shown below each plot. (E) Enlarged association plot for NW_026947487.1 (positions 63,652,500–77,760,726), with annotated genes; uncharacterized and predicted genes are not labeled. The coding variant identified by inheritance filtering of whole-genome sequencing data is shown in magenta.

**Table 2.** Scaffold name to placeholder number conversion chart for. **Figure 2A**. This table shows the conversion from reference genome assembly scaffold name to the placeholder number assigned for Figure 2A.

| **Scaffold Name** | **Placeholder Number on Figure 2A** |
|---|---|
| NW_026947858.1 | 1 |
| NW_026947855.1 | 2 |
| NW_026947844.1 | 3 |
| NW_026947716.1 | 4 |
| NW_026947715.1 | 5 |
| NW_026947660.1 | 6 |
| NW_026947616.1 | 7 |
| NW_026947606.1 | 8 |
| NW_026947603.1 | 9 |
| NW_026947567.1 | 10 |
| NW_026947537.1 | 11 |
| NW_026947536.1 | 12 |
| NW_026947535.1 | 13 |
| NW_026947534.1 | 14 |
| NW_026947533.1 | 15 |
| NW_026947532.1 | 16 |
| NW_026947530.1 | 17 |
| NW_026947529.1 | 18 |
| NW_026947528.1 | 19 |
| NW_026947526.1 | 20 |
| NW_026947525.1 | 21 |
| NW_026947524.1 | 22 |
| NW_026947523.1 | 23 |
| NW_026947522.1 | 24 |
| NW_026947521.1 | 25 |
| NW_026947520.1 | 26 |
| NW_026947518.1 | 27 |
| NW_026947517.1 | 28 |
| NW_026947516.1 | 29 |
| NW_026947515.1 | 30 |
| NW_026947514.1 | 31 |
| NW_026947513.1 | 32 |
| NW_026947512.1 | 33 |
| NW_026947511.1 | 34 |
| NW_026947510.1 | 35 |
| NW_026947509.1 | 36 |
| NW_026947508.1 | 37 |
| NW_026947507.1 | 38 |
| NW_026947506.1 | 39 |
| NW_026947505.1 | 40 |
| NW_026947504.1 | 41 |
| NW_026947503.1 | 42 |
| NW_026947502.1 | 43 |
| NW_026947501.1 | 44 |
| NW_026947500.1 | 45 |
| NW_026947499.1 | 46 |
| NW_026947498.1 | 47 |
| NW_026947497.1 | 48 |
| NW_026947496.1 | 49 |
| NW_026947495.1 | 50 |
| NW_026947494.1 | 51 |
| NW_026947493.1 | 52 |
| NW_026947492.1 | 53 |
| NW_026947491.1 | 54 |
| NW_026947490.1 | 55 |
| NW_026947489.1 | 56 |
| NW_026947488.1 | 57 |
| NW_026947487.1 | 58 |
| NW_026947486.1 | 59 |
| NW_026947485.1 | 60 |
| NW_026947484.1 | 61 |

Because the human exome-capture kit was designed to target coding regions, it is not expected to cover introns and may fail to capture guinea pig exons with insufficient sequence homology to human exons. Consequently, we found that 50% of coding bases were covered at a minimum depth of 1x, meaning low genotype call rates may exclude variants in associated regions from the GEMMA analysis. To address this, we directly assessed whole-genome sequencing data in the associated regions. We filtered variants according to the expected recessive inheritance pattern of long fur, requiring homozygosity for the alternate allele in all long-fur samples and either heterozygosity or homozygosity for the reference allele in all short-fur samples. Only the scaffold containing *FGF5* yielded variants consistent with this inheritance pattern (Table 3). Of the 11 variants identified, seven were intronic, three were in regions without annotated genes, and one was a coding variant, a C-to-T substitution in *FGF5* exon 2 (XM_003477669.5:c.403C>T; Figure 2E, magenta), which was also the most significant variant in the original GEMMA analysis (p = 4.27 × 10⁻¹³, β = 0.40, standard error = 0.04).

**Table 3.** Variant filtering for the long-fur trait. Variants in regions associated with long fur that follow the recessive inheritance pattern.

| **Scaffold** | **Position** | **Ref Allele** | **Alt Allele** | **Location** |
|---|---|---|---|---|
| NW_026947487.1 | 64181136 | A | G | no annotated genes |
| NW_026947487.1 | 65024971 | T | C | *CFAP299* intron |
| NW_026947487.1 | 65057798 | C | A | *CFAP299* intron |
| NW_026947487.1 | 65198870 | C | G,T | *CFAP299* intron |
| NW_026947487.1 | 65211707 | T | C | *CFAP299* intron |
| NW_026947487.1 | 65357151 | G | A | *FGF5* exon |
| NW_026947487.1 | 65489609 | C | T | no annotated genes |
| NW_026947487.1 | 65539295 | C | G | no annotated genes |
| NW_026947487.1 | 65600889 | A | T | *ANTXR2* intron |
| NW_026947487.1 | 65687333 | T | C | *ANTXR2* intron |
| NW_026947487.1 | 74308473 | GAGAAG | GAG | *CCSER1* intron |

### A premature stop codon in FGF5 confirms the molecular basis of recessive long fur

To independently validate the *FGF5* coding variant identified through genome-wide mapping and inheritance filtering, we performed Sanger sequencing of *FGF5* exon 2 in 26 samples, including 5 previously sequenced and 21 additional samples from the same breeders, pet shelters, and local pet owners. This panel included 11 long-fur and 15 short-fur animals. The same C-to-T substitution was confirmed in this panel (Figure 3A). In total, genotyping across all methods showed complete concordance between genotype and known phenotype: all long-fur samples were homozygous recessive for the substitution, whereas all short-fur samples carried at least one wild-type allele (Figure 3B, n = 70). No discordant genotype–phenotype pairs were observed, consistent with complete penetrance of the recessive long-fur phenotype.

**Figure 3.**
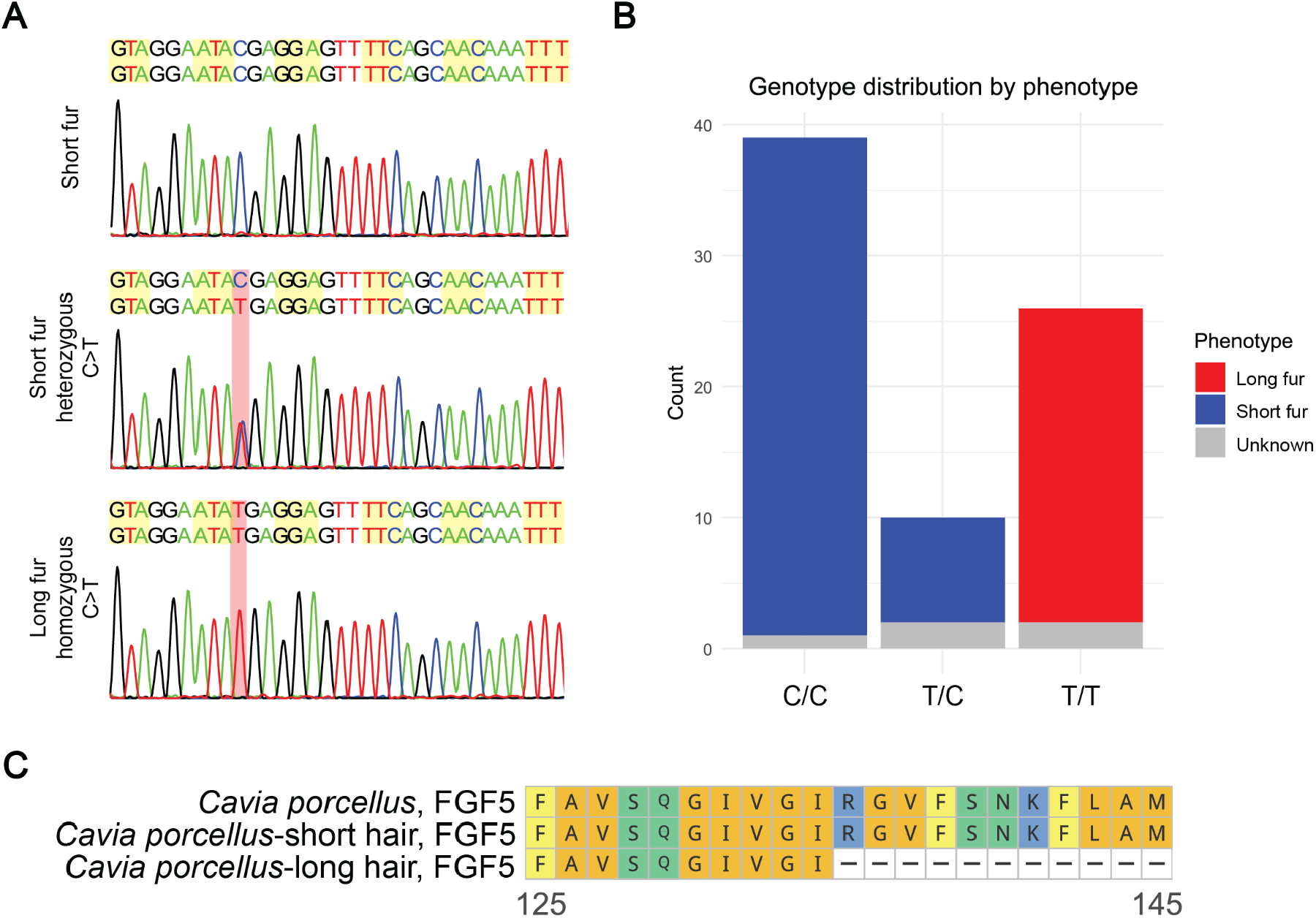
The associated *FGF5* variant introduces a premature stop codon. (A) Sanger sequencing chromatograms of *FGF5* exon 2 showing homozygous reference (top) and heterozygous (middle) short-fur genotypes and the homozygous C>T substitution in a long-fur sample (bottom). The variant is highlighted in red. In the heterozygous sample, the blue C trace is offset slightly to the right for visualization. Codons are shaded alternately yellow and white. (B) Genotype distributions by phenotype: short fur (blue), long fur (red), and unknown fur length (grey; pups). All samples with known phenotypes show genotype–phenotype concordance. (C) Alignment of the corresponding FGF5 protein region from the *Cavia porcellus* reference proteome and short-and long-fur samples. Amino acids are colored by chemical property. The long-fur genotype introduces a premature stop codon (p.Arg135*).

The C-to-T variant introduces a premature stop codon at amino acid 135 (H0UW37:p.Arg135*; Figure 3C). This variant was previously identified as a candidate for long fur through targeted sequencing of *FGF5* (Yu et al., 2018), and its recovery here through unbiased genome-wide mapping provides strong, independent genetic support for its association with long fur. Loss-of-function variants in *FGF5* have independently been identified as causes of long-hair phenotypes in several mammalian species, including cat, dog, mouse, hamster, rabbit, goat, and sheep (Carrion et al., 2024). In humans, *FGF5* mutations cause trichomegaly, characterized by excessively long eyelashes, further supporting a conserved role for FGF5 as a negative regulator of hair growth across mammals (Higgins et al., 2014). Together, these findings place the guinea pig long-fur variant within a previously-known spectrum of *FGF5* loss-of-function alleles associated with increased hair growth.

### Genetic mapping identifies a single locus containing *CELSR1* that is associated with dominant rough coat

The same genome-wide association approach and sequencing dataset used for long fur were applied to rough coat, treated as a binary trait based on the presence or absence of rosettes (rough coat = 28, smooth coat = 22, unknown = 4; Table 1, Figure S1). Samples were classified as having an unknown phenotype if the fur pattern was not sufficiently preserved for phenotypic assignment. As with the long-fur analysis, a centered kinship matrix was incorporated into the association model to account for relatedness, and GEMMA’s quality-control step restricted the analysis primarily to exonic variants, yielding 388,166 variants for association testing.

A single region on scaffold NW_026947523.1 exceeded the Bonferroni-corrected significance threshold (-log10(p) = 6.89) and was the only region to reach genome-wide significance (Figure 4A, Table 4), with no evidence of genomic inflation in the corresponding Q-Q plot (λ = 0.997; Figure 4B). Effect sizes (β) for the 39 significant variants indicated a 0.39-0.82 increase in the predicted probability of the rough-coat phenotype per additional effect allele (Figure 4C). LD was strong across all variants, ranging from r = 0.58 to 0.99. As expected, closely positioned variants showed stronger LD than more distant ones (Figure 4D).

**Figure 4.**
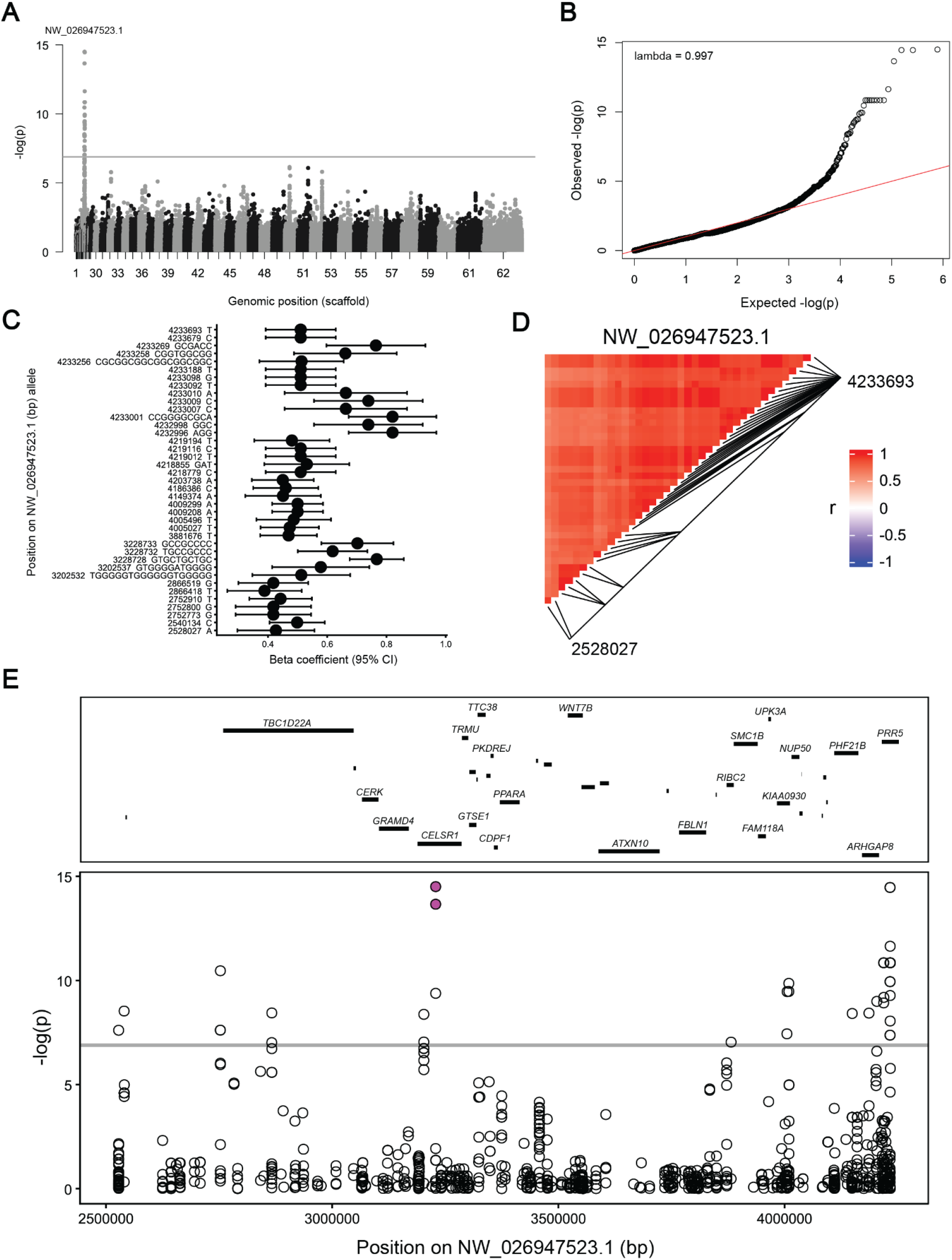
Genome-wide association mapping identifies a *CELSR1*-containing locus associated with rough coat. (A) Manhattan plot of genome-wide association results from GEMMA. Each point represents a variant, plotted by scaffold (numbered in descending order of size; full names are provided in Table 4) and association significance (−log₁₀(p)). The grey line indicates the Bonferroni-corrected significance threshold (−log₁₀(p) = 6.89). Significant variants occur only on scaffold 24 (NW_026947523.1). (B) Quantile–quantile (Q–Q) plot of observed versus expected −log₁₀(p) values (λ = 0.997). (C) β coefficients for significant variants, with scaffold positions (bp) and alleles indicated; bars represent 95% confidence intervals. (D) Linkage disequilibrium (LD) among significant variants on NW_026947523.1. Colors indicate pairwise correlation coefficients (*r*), from −1 (blue) through 0 (white) to 1 (red); variant positions (bp) are shown below the plot. (E) Enlarged association plot for NW_026947523.1 (positions 2,528,027–4,233,693), with annotated genes; uncharacterized and predicted genes are not labeled. The two coding variants identified by inheritance filtering of whole-genome sequencing data are shown in magenta.

**Table 4.**
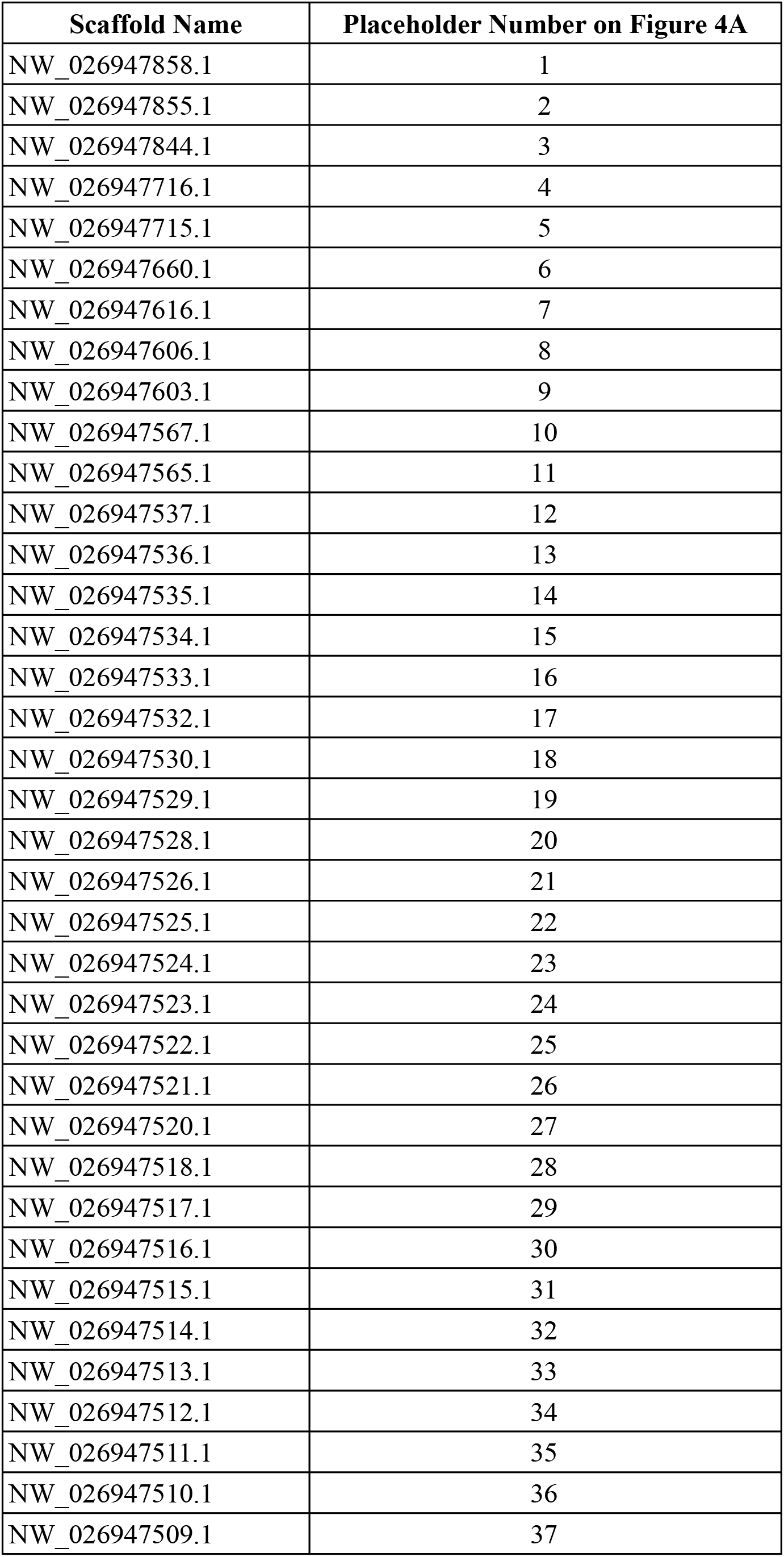
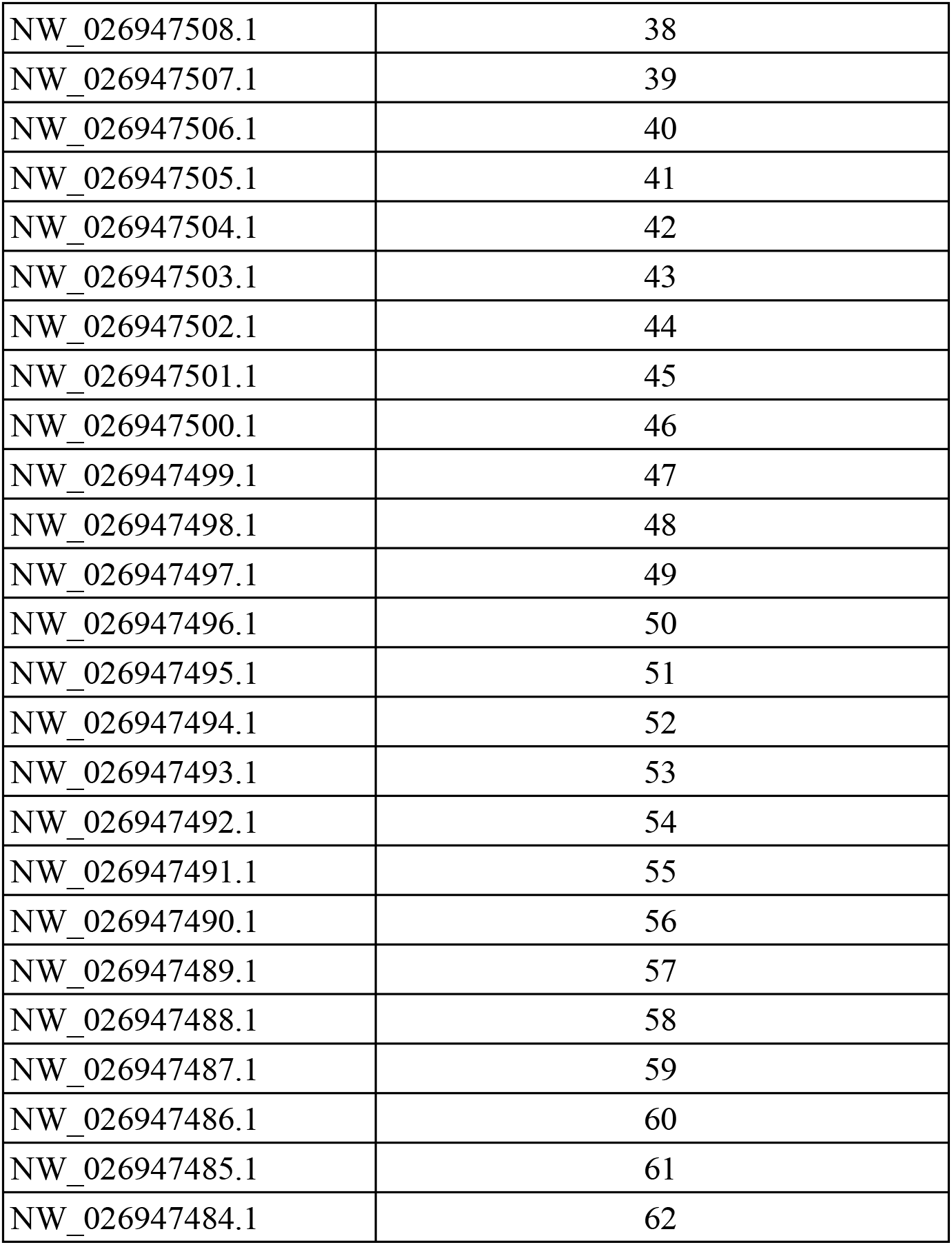
Scaffold name to placeholder number conversion chart for. **Figure 4A**. This table shows the conversion from reference genome assembly scaffold name to the placeholder number assigned for Figure 4A.

As with long fur, we directly assessed whole-genome sequencing data in the associated region to account for variants excluded from the GEMMA analysis. We then filtered variants in this region according to the expected dominant inheritance pattern of rough coat, requiring homozygosity for the reference allele in all smooth-coat samples and at least one copy of the alternate allele in all rough-coat samples (Table 5). Sixty-six variants passed this filter: 50 were intronic, 14 were in regions without annotated genes, and two were coding variants, both located in CELSR1 exon 2: XM_013158622.2:c.3841_3842insTGC (p = 3.12 × 10⁻¹⁵, β = 0.78, standard error = 0.05) and XM_013158622.2:c.3842_3843insGCC (p = 2.18 × 10⁻¹⁵, β = 0.62, standard error = 0.06) (Figure 4E, magenta dots).

**Table 5.**
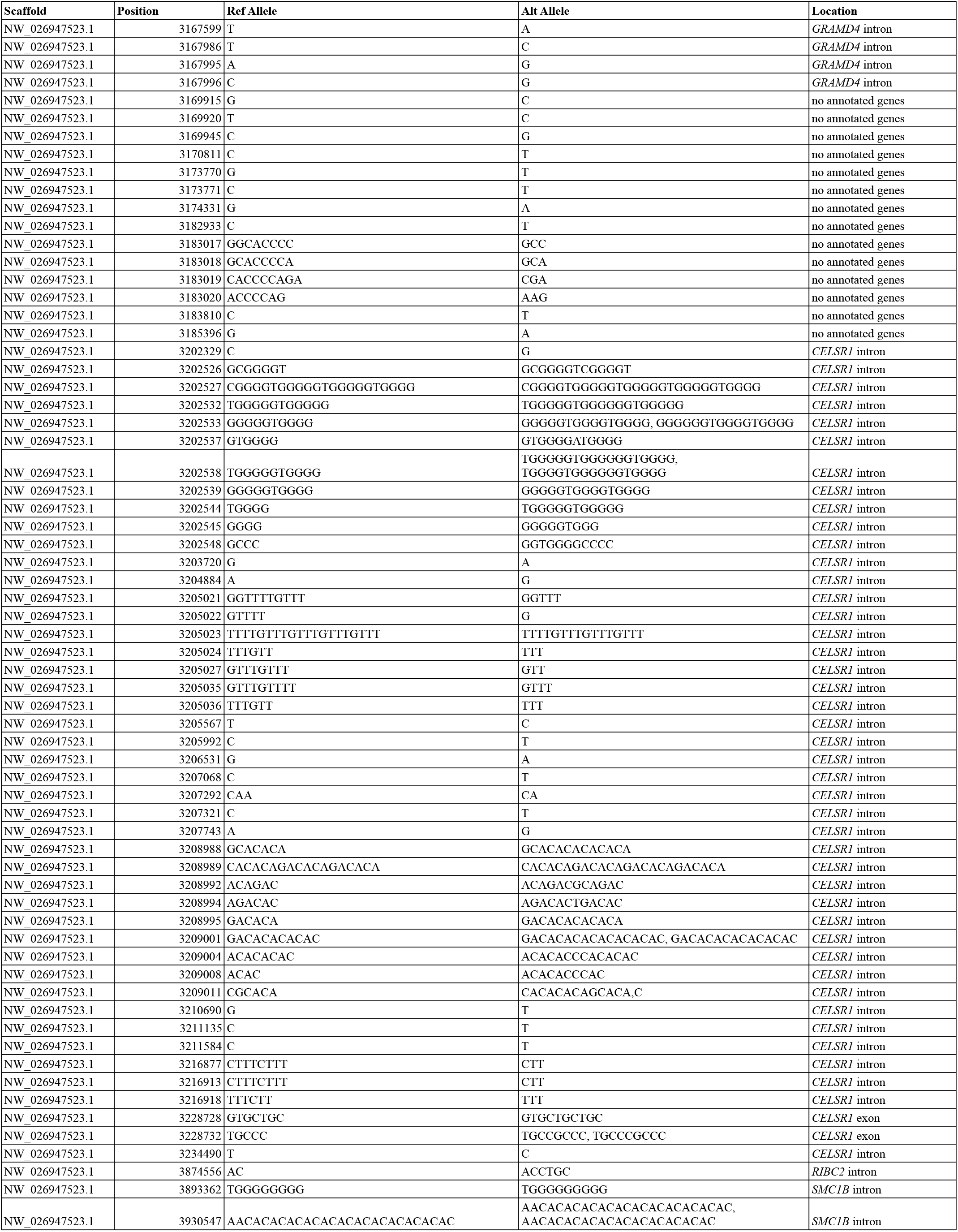
Variant filtering for the rough-coat trait. Variants in regions associated with rough coat that follow the dominant inheritance pattern.

### An in-frame proline insertion in CELSR1 segregates with dominant rough coat

To independently validate the *CELSR1* coding variants identified through genome-wide mapping and inheritance filtering, Sanger sequencing was performed across the candidate insertion sites in *CELSR1* exon 2 in a panel of 24 samples, including 9 previously sequenced samples and 15 additional samples; the panel comprised 11 rough-coat, 12 smooth-coat, and one unknown coat classification. One of the two three-base-pair insertions (XM_013158622.2:c.3841_3842insTGC) was not detected by Sanger sequencing in any sample. The other three-base-pair insertion, XM_013158622.2:c.3842_3843insGCC, was present in all rough-coat samples (Figure 5A). In total, genotyping across 65 samples with a known phenotype showed complete concordance between this insertion and coat phenotype: all smooth-coat samples were homozygous for the reference allele (*n* = 31), whereas all rough-coat samples carried at least one copy of the insertion, either as heterozygotes (*n* = 18) or homozygotes (*n* = 16) (Figure 5B). We observed no discordant genotype–phenotype pairs, consistent with the insertion being associated with dominant rough coat. At the protein level, the insertion adds a single proline residue immediately following an existing proline in the extracellular region of CELSR1 (H0VPZ8:p.Pro1194_Phe1195insPro; Figure 5C).

**Figure 5.**
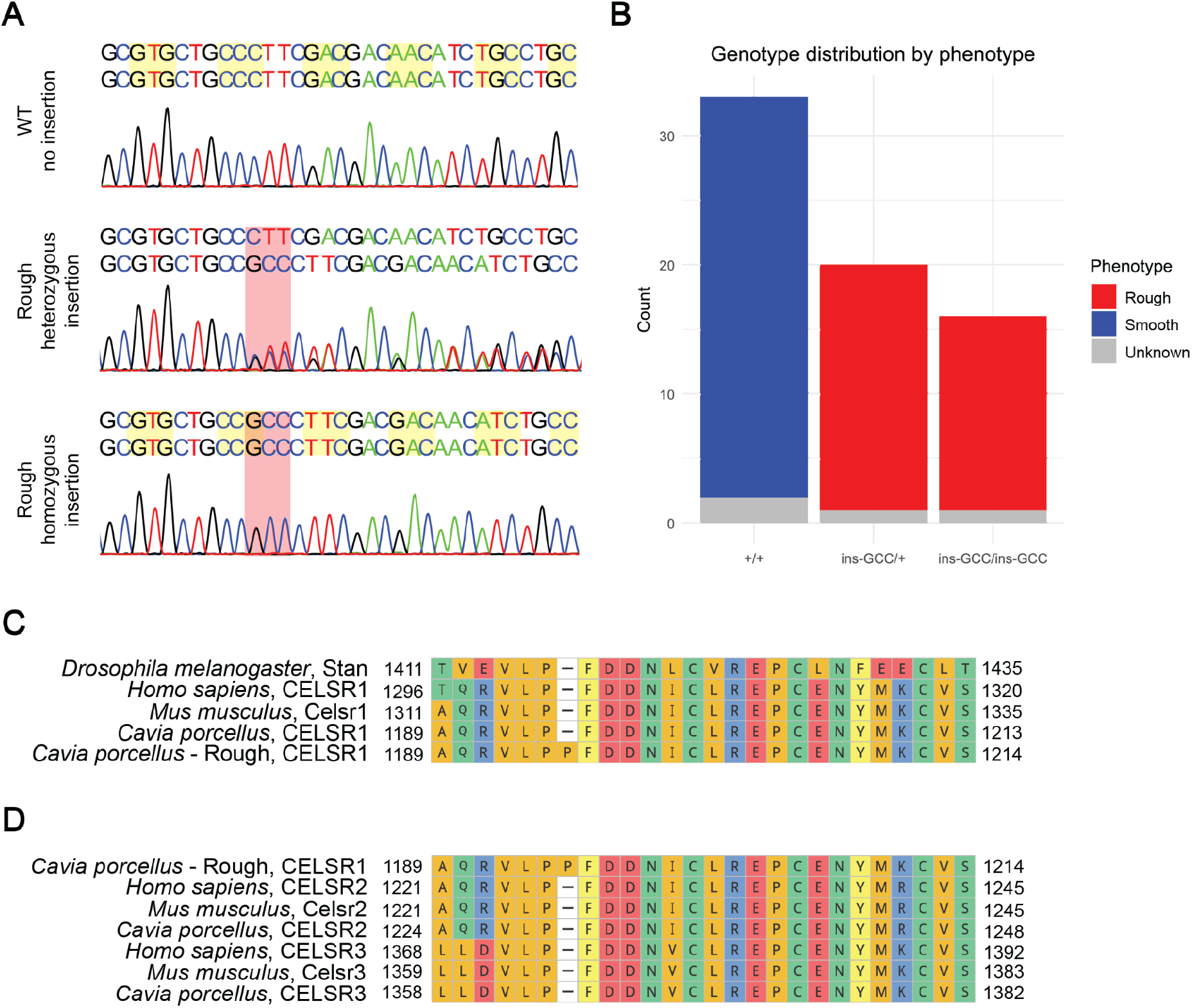
A GCC insertion introduces a proline into the extracellular region of CELSR1. (A) Sanger sequencing chromatograms of *CELSR1* exon 2 showing the homozygous reference genotype in a smooth-coat sample (top) and heterozygous and homozygous GCC insertion genotypes in rough-coat samples (middle and bottom; red highlights). Codons are shaded alternately yellow and white. (B) Genotype distributions by phenotype: smooth coat (blue), rough coat (red), and unknown phenotype (grey). All samples with known phenotypes show genotype–phenotype concordance. (C) Alignment of the corresponding CELSR1 region in *Drosophila melanogaster*, *Homo sapiens*, *Mus musculus*, and *Cavia porcellus* (reference proteome and rough-coat sample). The rough-coat sequence contains a proline insertion (p.Pro1194_Phe1195insPro). (D) Amino acid alignment of the corresponding region of CELSR1, CELSR2, and CELSR3 paralogs from *Homo sapiens*, *Mus musculus*, and *Cavia porcellus.* Amino acids are colored by chemical property (C-D).

To assess whether the insertion occurs within a conserved region of CELSR1, we examined sequence conservation at this locus across species. The sequence surrounding the insertion site is highly conserved among CELSR1 orthologs, with strong conservation observed among guinea pig (CELSR1), mouse (Celsr1), human (CELSR1), and *Drosophila* (Stan/Flamingo) (Figure 5C). The corresponding region is also highly similar among the three mammalian CELSR/Celsr paralogs in guinea pig, mouse, and human (Figure 5D). Thus, the insertion occurs within a region of CELSR1 that is highly conserved across species and shared among mammalian CELSR/Celsr paralogs.

### The Celsr1 rough-coat variant localizes normally to cell-cell junctions and recruits core PCP components

Celsr1 is an atypical cadherin that localizes to cell-cell junctions through homophilic interactions with Celsr1 on the opposing cell (Figure 6A). Structural studies of mouse Celsr1 and human CELSR1 indicate that rough-coat insertion occurs at the C-terminus of the Flamingo box (Fbox) in the extracellular domain (Figure 6B, S2) (Bandekar et al., 2025; De-la-Torre et al., 2018; Tamilselvan & Sotomayor, 2024). The Fbox is structurally similar to the membrane-adjacent domain (MAD) of protocadherins, which mediates cis interactions that stabilize cadherin adhesion in trans between neighboring cells (Bandekar et al., 2025; De-la-Torre et al., 2018; Tamilselvan & Sotomayor, 2024). This raises the possibility that the insertion disrupts cis-clustering similarly to *Celsr1^Crsh^*, a *Celsr1* missense variant that impairs cis-clustering and, as a result, fails to enrich normally at cell-cell junctions (Stahley et al., 2021). To test this, we examined junctional localization of the rough-associated variant.

**Figure 6.**
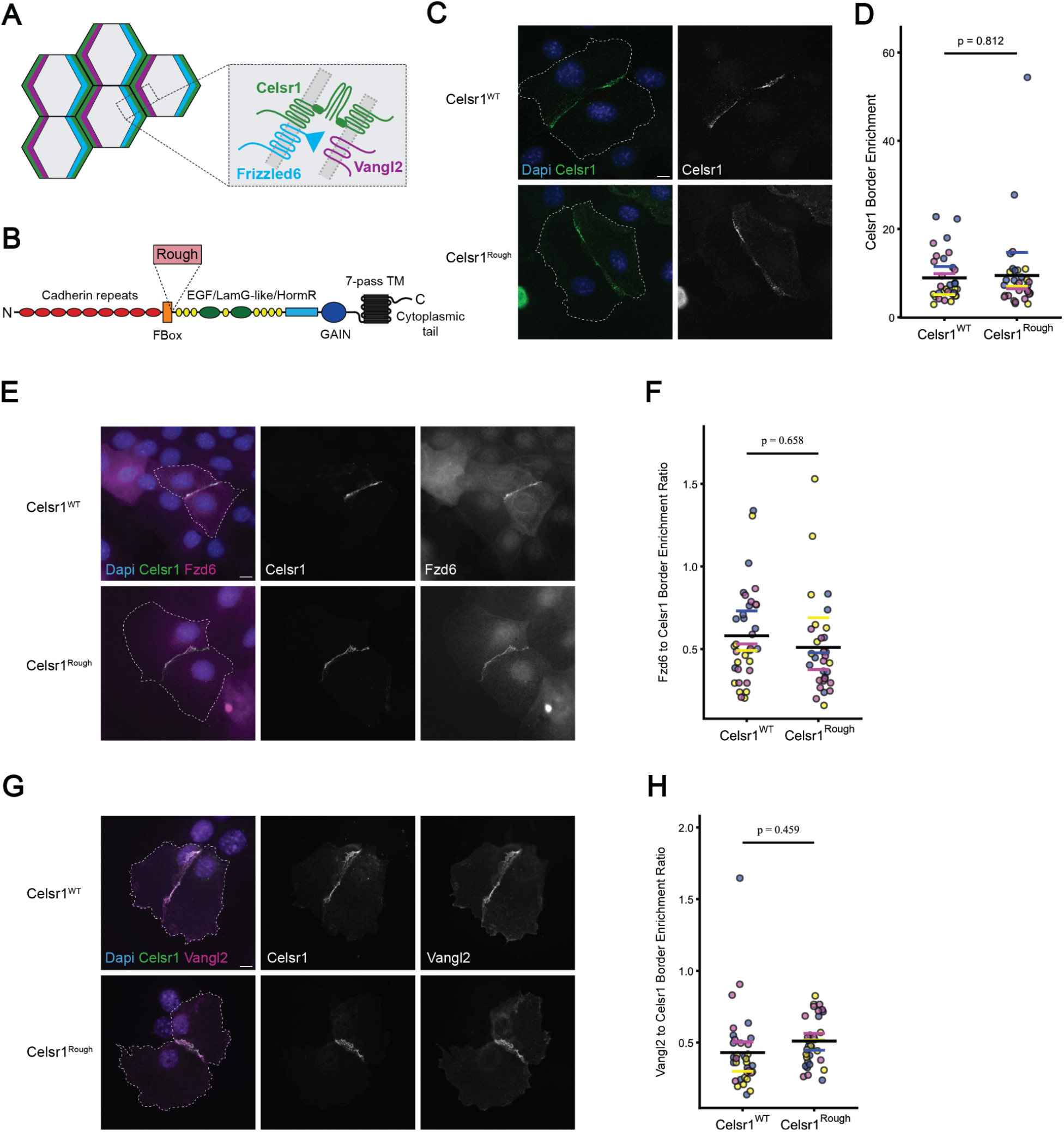
Celsr1^Rough^ localizes to cell-cell junctions and recruits PCP proteins. (A) Schematic of core PCP proteins: Celsr1 (green), Fzd6 (blue), and Vangl2 (magenta), showing their localization and extracellular interactions at an asymmetric junction. (B) Celsr1 domain structure, with cadherin repeats 1–9 (red), Fbox (orange), EGF-like domains 1–8 (yellow), Laminin G-like domains 1 and 2 (green), HormR (light blue), GAIN (dark blue), and the seven-pass transmembrane domain (black rectangles). The Rough insertion is located at the C-terminus of the FBox. (C) Representative images of mouse keratinocytes expressing Celsr1^WT^-GFP (top) or Celsr1^Rough^-GFP (bottom). (D) Quantification of Celsr1-GFP border enrichment (WT, n = 32; Rough, n = 33; two-tailed paired *t*-test, p = 0.812). (E) Representative images of cells coexpressing Fzd6-tdTomato and Celsr1^WT^-GFP (top) or Celsr1^Rough^-GFP (bottom). (F) Ratio of Fzd6 to Celsr1 border enrichment (WT, n = 34; Rough, n = 32; two-tailed paired *t*-test, p = 0.658). (G) Representative images of cells coexpressing 3×FLAG-Vangl2 and Celsr1^WT^-GFP (top) or Celsr1^Rough^-GFP (bottom). (H) Ratio of Vangl2 to Celsr1 border enrichment (WT, n = 34; Rough, n = 32; two-tailed paired *t*-test, p = 0.459). White dashed lines indicate cell outlines; scale bar, 10 μm (C, E, G). In (D, F, H), cell pairs from three independent experiments are colored blue, yellow, and magenta; colored bars indicate experiment means, and black bars indicate the overall mean.

We transfected mouse keratinocytes, a well-established system for studying planar cell polarity *in vitro* (Devenport & Fuchs, 2008), with GFP-tagged constructs encoding the rough-coat insertion at the corresponding position in the mouse *Celsr1* sequence (*Celsr1^Rough^*). To assess junctional localization, we expressed Celsr1^WT^ and Celsr1^Rough^ separately, so the junctions we analyzed formed between cells expressing the same Celsr1 construct. We evaluated protein enrichment at cell-cell junctions using a junction enrichment assay, which measures the ratio of the fluorescence signal at the junction to the signal along the rest of the cell border (Devenport & Fuchs, 2008). Junction enrichment did not differ between Celsr1^WT^ and Celsr1^Rough^ (p = 0.812) (Figure 6C-D). This indicates that the rough-coat insertion does not impair Celsr1 junctional localization, arguing against a comparable disruption of cis-clustering.

*In vivo*, Celsr1 recruits Fzd6 and Vangl2 to opposite sides of the cell-cell junction, forming an asymmetric complex (Figure 6A). In the *in vitro* system, Fzd6 and Vangl2 require Celsr1 for junctional localization (Devenport & Fuchs, 2008), allowing their recruitment to serve as a readout for whether the rough-coat variant alters Celsr1 interactions with the core PCP proteins. We coexpressed tagged versions of Fzd6 with either Celsr1^WT^ or Celsr1^Rough^ and then calculated the ratio of Fzd6 border enrichment to the corresponding Celsr1 border enrichment, as described above. The same experiment was performed with Vangl2. Both Celsr1^WT^ and Celsr1^Rough^ recruited Fzd6 and Vangl2 to cell-cell junctions at comparable levels (Fzd6 p = 0.658, Vangl2 p = 0.459) (Figure 6E-H), demonstrating that the rough-coat insertion preserves the ability of Celsr1 to recruit these core PCP components.

Together, these findings establish that Celsr1^Rough^ retains key molecular properties of Celsr1, including junctional localization and recruitment of the core PCP proteins Fzd6 and Vangl2. The fact that these properties are retained indicates that Celsr1^Rough^ does not act through a simple loss-of-function mechanism, distinguishing it from previously characterized *Celsr1* alleles.

## DISCUSSION

Using genome-wide mapping, this study identifies the likely molecular basis of two classical guinea pig coat phenotypes described more than a century ago. Mapping of long fur identified a premature stop codon in *FGF5*, providing independent genetic support for a previously proposed candidate variant (Yu et al., 2018). Mapping of rough coat identified an in-frame proline insertion in the extracellular domain of the core PCP protein CELSR1. This variant is associated with a fully penetrant, dominant hair-patterning phenotype, distinguishing it from previously characterized *Celsr1* alleles in mice, which produce comparable coat phenotypes only in homozygotes (Basta et al., 2022; Ravni et al., 2009). The dominant inheritance of rough coat, along with preserved junctional localization and recruitment of core PCP components in cultured cells, suggests that the insertion alters CELSR1 function in a manner distinct from simple loss of function.

If the CELSR1 insertion acted as a simple loss-of-function allele, dominance would require haploinsufficiency; however, heterozygous carriers of *Celsr1* loss-of-function alleles in mice show no coat phenotype, arguing against this model (Basta et al., 2022; Ravni et al., 2009). The rough-coat variant is similarly distinct from *Celsr1^Crsh^*, which carries a missense mutation between cadherin repeats 7 and 8 that disrupts the cis-interactions underlying Celsr1 cluster formation and junctional enrichment (Curtin et al., 2003; Stahley et al., 2021). Celsr1^Crsh^ may act as a partial dominant negative by limiting cluster size (Stahley et al., 2021), yet wild-type Celsr1 could still cluster sufficiently in heterozygotes, consistent with a lack of hair orientation defects. In contrast, the rough-coat variant is fully dominant despite normal junctional localization, suggesting a more effective disruption of wild-type Celsr1 function in heterozygotes. In *Drosophila*, Frizzled binding is proposed to induce a conformational change in Flamingo that enables its trans-homophilic interaction with Flamingo on the neighboring cell, which in turn promotes asymmetric recruitment of Strabismus, the *Drosophila* ortholog of Vangl, on that side of the junction (Strutt et al., 2023). If the rough-coat insertion alters a conformational change required for this process while retaining the ability to cis- and trans-associate with wild-type CELSR1, it could incorporate directly into PCP complexes alongside wild-type protein and disrupt the conformational relay from within, providing a basis for its dominance.

Although the mechanism by which the insertion alters CELSR1 function remains unresolved, the range of hair-patterning phenotypes in mouse PCP mutants provides a framework for understanding rough coat. The most severe coat phenotype, resembling full rough, results from epidermal loss of both *Vangl1* and *Vangl2*, which eliminates redundancy between these core PCP components (Cetera et al., 2017). Homozygous *Celsr1* null mice show a similar but generally milder, variably penetrant phenotype (Basta et al., 2022; Ravni et al., 2009). In contrast, epidermal-specific loss of *Vangl2* or loss of *Fzd6* produces a milder, paw-restricted phenotype in adults, consistent with residual PCP activity due to compensation by Vangl1 or Fzd3, respectively (Cetera et al., 2017; H. Chang et al., 2016; Dong et al., 2018). Together, these mouse models support a spectrum of PCP activity in which the extent of hair-orientation defects reflects the degree of pathway disruption: follicle orientation on the paw is the most sensitive readout of reduced PCP activity, while full-body whorls reflect near-complete elimination of pathway function. Notably, the study that identified *Fzd6* as the first mammalian PCP gene shown to control hair orientation drew comparison to Wright’s whorled guinea pigs (Guo et al., 2004).

Rough-coat guinea pigs exhibit a similar range of phenotypic severity, from rosettes distributed across the body to altered fur orientation restricted to the paws. Wright’s breeding studies established that an unlinked, semidominant modifier influences this variation (Wright, 1949). The modifying allele *M* suppresses whorls while *m* permits their spread across the body. Because *M* is present in wild *Cavia* species, it is likely the ancestral state, with *m* arising through a subsequent reduction or loss of PCP activity. With this framing, paw-restricted fur misorientation likely represents the baseline effect of the *CELSR1* insertion, and *m* can be viewed as enhancing the rough phenotype. Mice provide a precedent for a naturally occurring modifier of a PCP hair phenotype, in which a spontaneous, recessive deletion in *Astn2* enhances the hair orientation defects of *Fzd6* null animals (H. Chang et al., 2015). The ancestral state of the modifier is also relevant to the interpretation of the *in vitro* results. Since wild-type mouse keratinocytes presumably lack the enhancing effect of *m*, the assays assess the baseline defect that only impacts fur on the paws, which may not be disruptive enough to detect *in vitro*.

The dependence of PCP phenotypes on the extent of pathway disruption also raises the question of why the *CELSR1* insertion is associated with a coat phenotype without neural tube defects or other developmental abnormalities. If the rough-coat variant acts by incorporating into and disrupting wild-type complexes, as proposed above, its effects may depend on how PCP complexes are organized and maintained in different tissues. One possibility is that skin patterning depends more heavily on stable, long-lived junctions between relatively static cells, whereas neural tube closure involves extensive cell rearrangements that may be less dependent on long-term, complex stability (Basta et al., 2021; Moran & Zohn, 2026). A second possibility is that redundancy among CELSR paralogs differs by tissue and/or species: although Celsr1 is largely non-redundant with Celsr2 in mouse skin (Basta et al., 2022), CELSR2 or CELSR3 could contribute more substantially to neural tube closure in the guinea pig, buffering the variant’s effects in this tissue.

Several limitations of this study should be noted. The *in vitro* assays used here were restricted to junctions between isolated cell pairs and lacked the super-resolution imaging needed to assess their asymmetry. Because the orientation of PCP junction asymmetry is uniform across entire tissues, the insertion could potentially disrupt the coordination of PCP asymmetry across a multicellular field, which these assays cannot detect. In addition, these experiments were conducted using the rough-coat insertion introduced into the mouse *Celsr1* sequence rather than its native guinea pig context, which may not fully recapitulate the variant’s behavior. Finally, the identity of Wright’s modifier remains unknown, and confirming the mechanisms proposed here will require its identification through future genetic mapping.

More broadly, these findings highlight the continued power of forward genetics to reveal developmental mechanisms from natural variation. In hindsight, Wright’s breeding studies were an early investigation of variation in a PCP-dependent phenotype, conducted decades before PCP genes were identified in *Drosophila* (Chae et al., 1999; Gubb & García-Bellido, 1982; Taylor et al., 1998; Usui et al., 1999; Wolff & Rubin, 1998). Naturally occurring alleles, such as the dominant CELSR1 variant described here, can reveal functional properties of a pathway that engineered loss-of-function models are not designed to capture. Together, these findings show how modern genomics can resolve the molecular basis of PCP phenotypes first defined by classical breeding studies a century ago.

## MATERIALS AND METHODS

### Key Resources (Table 6)

| Reagent or resource type | Name | Reference/ source | Identifier | Additional information |
|---|---|---|---|---|
| Software/packages | Trim Galore | (Krueger, 2023) |  |  |
| Software/packages | BWA-MEM | (Li, 2013) |  |  |
| Software/packages | SAMtools/BCFtools | (Danecek et al., 2021) |  |  |
| Software/packages | Integrative Genomics Viewer | (Robinson et al., 2011) |  |  |
| Software/packages | GEMMA | (X. Zhou & Stephens, 2012) |  |  |
| Software/packages | RStudio package: qqman | (Turner, 2018) |  |  |
| Software/packages | RStudio package: locuszoomr | (Boughton et al., 2021) |  |  |
| Software/packages | PLINK 1.9 | (C. C. Chang et al., 2015) |  |  |
| Software/packages | RStudio package: LDheatmap | (Shin et al., 2006) |  |  |
| Software/packages | RStudio package: ComplexHeatmap | (Gu et al., 2016) |  |  |
| Software/packages | RStudio package: sangerseqR | (Hill et al., 2014) |  |  |
| Software/packages | RStudio package: ggmsa | (L. Zhou et al., 2022) |  |  |
| Plasmid | pEGFPN1-Celsr1 | (Devenport & Fuchs, 2008) |  |  |
| Plasmid | pEGFPN1-Celsr1-Rough | This study |  |  |
| Plasmid | K14-HA-Fzd6-tdTom | (Cetera et al., 2023) |  |  |
| Plasmid | K14-FLAG-Vangl2 | Gift from Danelle Devenport |  |  |
| Antibody | Anti-GFP (Chicken, polyclonal) | Abcam | Catalog # ab13970 | 1:2000 |
| Antibody | Anti-RFP (Rabbit, polyclonal) | Rockland | Catalog # 600-401-379 | 1:200 |
| Antibody | Anti-DYKDDDDK (Goat, polyclonal) | Bethyl Laboratories | Catalog # A190-101A | 1:400 |
| Antibody | Anti-chicken, Alexa Fluor 488 (Goat) | Invitrogen | Catalog # A-11039 | 1:1500 |
| Antibody | Anti-chicken Alexa Fluor 488 (Donkey) | Invitrogen | Catalog # A78948 | 1:2000 |
| Antibody | Anti-rabbit, Alexa Fluor 555 (Donkey) | Invitrogen | Catalog # A-31572 | 1:1500 |
| Antibody | Anti-goat, Alexa Fluor 647 (Donkey) | Invitrogen | Catalog # A-21447 | 1:1500 |

**Table 6.** Key resources table. Indicates resources used for this research.

| Reagent or resource type | Name | Reference/source | Identifier | Additional information |
|---|---|---|---|---|
| Software/packages | Trim Galore | (Krueger, 2023) |  |  |
| Software/packages | BWA-MEM | (Li, 2013) |  |  |
| Software/packages | SAMtools/BCFtools | (Danecek et al., 2021) |  |  |
| Software/packages | Integrative Genomics Viewer | (Robinson et al., 2011) |  |  |
| Software/packages | GEMMA | (X. Zhou & Stephens, 2012) |  |  |
| Software/packages | RStudio package: qqman | (Turner, 2018) |  |  |
| Software/packages | RStudio package: locuszoomr | (Boughton et al., 2021) |  |  |
| Software/packages | PLINK 1.9 | (C. C. Chang et al., 2015) |  |  |
| Software/packages | RStudio package: LDheatmap | (Shin et al., 2006) |  |  |
| Software/packages | RStudio package: ComplexHeatmap | (Gu et al., 2016) |  |  |
| Software/packages | RStudio package: sangerseqR | (Hill et al., 2014) |  |  |
| Software/packages | RStudio package: ggmsa | (L. Zhou et al., 2022) |  |  |
| Plasmid | pEGFPN1-Celsr1 | (Devenport & Fuchs, 2008) |  |  |
| Plasmid | pEGFPN1-Celsr1-Rough | This study |  |  |
| Plasmid | K14-HA-Fzd6-tdTom | (Cetera et al., 2023) |  |  |
| Plasmid | K14-FLAG-Vangl2 | Gift from Danelle Devenport |  |  |
| Antibody | Anti-GFP (Chicken, polyclonal) | Abcam | Catalog # ab13970 | 1:2000 |
| Antibody | Anti-RFP (Rabbit, polyclonal) | Rockland | Catalog # 600-401-379 | 1:200 |
| Antibody | Anti-DYKDDDDK (Goat, polyclonal) | Bethyl Laboratories | Catalog # A190-101A | 1:400 |
| Antibody | Anti-chicken, Alexa Fluor 488 (Goat) | Invitrogen | Catalog # A-11039 | 1:1500 |
| Antibody | Anti-chicken Alexa Fluor 488 (Donkey) | Invitrogen | Catalog # A78948 | 1:2000 |
| Antibody | Anti-rabbit, Alexa Fluor 555 (Donkey) | Invitrogen | Catalog # A-31572 | 1:1500 |
| Antibody | Anti-goat, Alexa Fluor 647 (Donkey) | Invitrogen | Catalog # A-21447 | 1:1500 |

### Animals and sample collection

All procedures involving animals in this study were approved by the University of Minnesota’s Institutional Animal Care and Use Committee (IACUC; protocols 2106-39180A, 2403-41891A). Full phenotype, genotype, and sequencing information for all samples are listed in Table 1. Ear clips were taken from frozen carcasses of previously deceased guinea pigs donated by breeders and pet owners in the Minneapolis, Minnesota area. Animals were selected based on phenotype and sample availability. Phenotypes were assessed by fur pattern, photographs, and owner-provided information.

### Sample sequencing and data processing

Genomic DNA was extracted from tissue using a QIAamp DNA Mini Kit. Thirty-two high molecular weight DNA samples were sequenced using the Illumina platform in collaboration with the University of Minnesota Genomics Center (UMGC). Library prep was performed with the Illumina DNA Prep with Exome 2.5 Enrichment, (S) Tagmentation Set B kit, which is a human whole-exome enrichment kit. A human exome kit was used because commercial guinea pig kits were not available. Samples were sequenced at high coverage (expected >30x for human samples) using a 150bp PE NovaSeq SPrime single lane. Exons containing variants that correlate with the phenotypes of interest were captured by this method (Figure S3). Two high molecular weight samples were prepared for whole genome sequencing with the Illumina platform at UMGC. Nextera DNA Flex (now called Illumina DNA Prep) was used for library preparation and samples were sequenced at high coverage (expected 30x) using a 150bp PE NovaSeq S4 single lane. Twenty high molecular weight DNA samples were prepared for whole genome sequencing using the Ultima platform at UMGC. Samples were sequenced at high coverage (expected ∼33x), using NEB NExt PCR free library prep kit with UG100 adapters and UG100 Solaris Free sequencing chemistry.

Adapters were trimmed from sequencing data using Trim Galore 0.6.10 with the –paired option for Illumina sequenced samples (Krueger, 2023). Trimmed data was then aligned to mCavPor4.1 (NCBI RefSeq accession GCF_034190915.1) using BWA-MEM and SAMtools, filtering for reads with a map quality of 30 or higher (Danecek et al., 2021; Li, 2013). This is a scaffold-level genome assembly. Bam files were sorted and indexed using SAMtools, and variants were called using BCFtools mpileup and call (Danecek et al., 2021). SAMtools was used for the following analyses (Danecek et al., 2021). Percentage of reads mapped confidently for exome-captured samples was determined by the number of primary reads mapped with a mapping quality of ≥ 30, using the flagstat command, compared to the total number of reads present in fastq files. Genome coverage for exome-captured samples was determined using the coverage command, where the added average covbases between samples for each scaffold was divided by the added endpos for each scaffold. Exome coverage was determined for each sample by taking the number of positions with ≥1 read using the depth command with the -b argument with a bed file for locations annotated as cds in the mCavPor4.1 reference genome gtf file. This value was then divided by the total positions in the cds as annotated by the mCavPor4.1 reference genome gtf file and averaged over all exome-captured samples. Mean depth for exome-captured samples was calculated using the depth command at locations annotated as cds in the mCavPor4.1 reference genome gtf file, averaged for each sample to get mean depth per sample, and then averaged over all exome-captured samples. Mean depth for whole-genome sequencing samples was calculated using the coverage command and weighted by scaffold length [(sum for all samples and scaffolds (mean depth per scaffold x scaffold length))/(summed scaffold lengths x number of samples)]. Coverage was also assessed using Integrative Genomics Viewer (IGV) (Robinson et al., 2011).

### Genotype-phenotype association

A univariate linear mixed model (LMM) from the program GEMMA 0.98.5 was used to identify variants associated with the long-fur and rough-coat phenotype in guinea pigs (X. Zhou & Stephens, 2012). The phenotypes were treated as categorical (present, absent, or unknown) according to Table 1. Long-fur phenotype was treated as unknown when the sample was a pup without fully grown-out fur, and the rough-coat phenotype was treated as unknown if the fur pattern was not sufficiently preserved for phenotypic assignment. A centered relatedness matrix was calculated by GEMMA and then used in the LMM to adjust association scores for relatedness. The likelihood ratio test was used to test significance. For quality control, GEMMA ignores samples marked as unknown phenotype and excludes variants where >5% of the genotypes are missing. Because exons with low homology to the human exome capture probes were poorly covered, some scaffolds had no variants passing GEMMA’s quality-control filters and were therefore absent from the dataset entirely. As a result, the remaining scaffolds no longer formed a consecutive numerical series; for plotting purposes, each remaining scaffold was assigned a consecutive placeholder number, with the conversion between placeholder number and original scaffold name recorded for reference (Table 2 and Table 4). Significance values (-log_10_ p-value) were graphed in the Manhattan plot by location in the genome according to scaffold. The significance threshold was determined using the Bonferroni correction (0.05/386,066 = 6.89 for long-fur analysis and 0.05/388,166 = 6.89 for rough-coat analysis). The RStudio packages qqman and locuszoomr were used to create Manhattan plots (Boughton et al., 2021; Turner, 2018).

Because the human exome kit did not capture all guinea pig exons, whole-genome sequencing data in the regions of interest were used to identify additional variants consistent with the expected inheritance pattern for each trait. Because whole-genome sequencing is expected to cover the genome, variants missing genotype calls in more than 5% of whole-genome-sequenced samples were excluded from this filtering step.

### Quantile-Quantile plot

Expected and observed significance values were compared for GEMMA results using RStudio package qqman (Turner, 2018). Significance values were converted to chi-square test statistics using a chi-square distribution with one degree of freedom, and lambda (λ) was calculated as the ratio of the median observed chi-square statistic to the expected median of the chi-square distribution with one degree of freedom.

### Beta Coefficient forest plot

The effect size (β) and standard error were used from the GEMMA results to calculate the 95% confidence intervals (β ⼟ 1.96×SE) of variants rising above the Bonferroni correction significance threshold. If the effect size was negative, the opposing allele was used for analysis, and alleles used are listed in the plot.

### Linkage disequilibrium (LD) plot

PLINK 1.9 (C. C. Chang et al., 2015) was used to calculate the r values between significant variants (the same alleles from the Beta Coefficient forest plot). R values were then graphed in a linkage disequilibrium (LD) plot using RStudio package LDheatmap and ComplexHeatmap (Gu et al., 2016). R values were not graphed for scaffolds with a single significant variant.

### Sanger sequencing

PCR amplification was used to amplify the variant and surrounding DNA using the primer sets in Table 7. The Zymo Research DNA Clean and Concentrator kit was used before sending the samples to GENEWIZ for Sanger sequencing. Trace files were analyzed using SnapGene and compared with phenotypes from Table 1. The RStudio package sangerseqR was used to create trace file comparisons (Hill et al., 2014). The region of interest for *FGF5* was sequenced on 11 long-fur and 15 short-fur samples and the *CELSR1* region of interest was sequenced on 11 rough and 12 smooth samples, as well as one sample with an unknown phenotype.

**Table 7.**
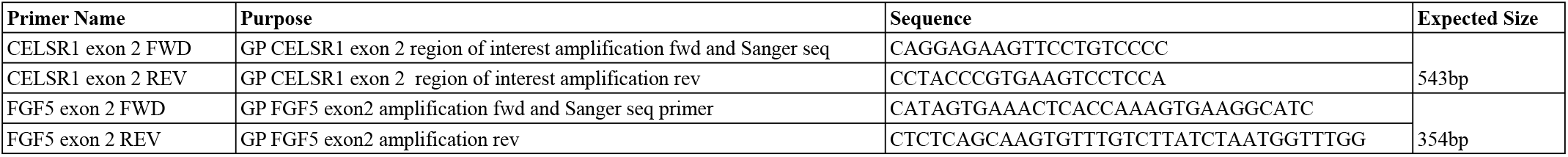
Oligonucleotides table. Indicates oligonucleotides used for Sanger sequencing.

### Protein comparison

Consensus sequences for samples were pulled from Bam files using BCFtools consensus and compared to Uniprot protein sequences for FGF5 and CELSR1-CELSR3 (Danecek et al., 2021). The Uniprot accession H0UW37 was used for FGF5. The following accessions were used for CELSR1/Celsr1-CELSR3/Celsr3: Q9V5N8, Q9NYQ6, O35161, H0VPZ8, Q9HCU4, Q9R0M0, A0A286Y5A0, Q9NYQ7, Q91ZI0, and A0A286XSG3. The RStudio package ggmsa was used to create protein comparisons (L. Zhou et al., 2022).

### Cells and cell lines

CD1 mouse keratinocytes were derived from P0 dorsal skins and cultured in E-media supplemented with 15% fetal bovine serum and 0.05 mM Ca^2+^ at 37°C with 5% CO_2_. For experiments, passage 14-20 cells were cultured on fibronectin-treated coverslips and transfected with constructs using the Effectene transfection reagent (Qiagen). Cells were switched to 1.5 mM Ca^2+^ high-calcium media, 16 hours post-transfection, and fixed in 4% paraformaldehyde for 10 minutes, 24 hours post-media change.

### Plasmid constructs

Plasmids used in this study are listed in Table 6. For this study, pEGFPN1-Celsr1^Rough^ (Celsr1^Rough^-GFP) construct was generated by VectorBuilder from the pEGFPN1-Celsr1^WT^ (Celsr1^WT^-GFP) construct (Devenport & Fuchs, 2008).

### Immunostaining

Fixed cells were washed in PBS and permeabilized in 0.1% Triton X-100 in PBS (PBT). Cells were then incubated with primary antibodies in PBT for one hour, washed with PBT, incubated with secondary antibodies in PBT for 30 minutes, washed with DAPI (0.1μg/mL), washed with PBT, and then mounted with ProLong Gold Antifade Mountant (Invitrogen). Antibody dilutions are listed in Table 6.

### Microscopy

Cells were imaged on a Nikon Ti-2 wide-field system. Positions were chosen using the GFP signal at 20x magnification and then imaged at 40x using an oil immersion objective. Cells were imaged using auto-exposure due to differences in plasmid expression between cell pairs. To be included in the analysis, both cells in the pair had to express all transfected constructs and could not share a cell-cell border with more than one cell expressing Celsr1-GFP.

### Border enrichment calculations and statistical analysis

Border enrichment was measured using the freehand line tool in ImageJ. For each cell pair, average fluorescence intensity was measured along the shared cell-cell border, along the non-shared borders, and for the background. The background value was removed from all measurements, and the border enrichment was calculated by the ratio of the average shared border fluorescence to the average non-shared border fluorescence. For Fzd6 and Vangl2 transfection experiments, the ratio of Fzd6 or Vangl2 border enrichment to Celsr1 border enrichment was calculated.

At least 10 cell pairs were analyzed for each experiment, and three replicates were performed for each condition. The average border enrichment values across all cell pairs for an experiment were used in a paired two-tailed *t*-test for statistical analysis.

## DATA ACCESSIBILITY

Sequencing data will be available through NCBI SRA upon acceptance.

## ACKNOWLEDGEMENTS

We thank Martha Panda and Paul Weinand for donations of samples and images, as well as for sharing their insights about rough-coat guinea pigs. We also thank the pet shelters and foster programs throughout the Twin Cities that contributed samples. We thank Julaine Roffers-Agarwal, Evalyn Beall, Connie Corcoran, and Thu Nguyen for their contributions to sample collection during pilot studies. We thank members of the Cetera, Isabella, Artinger, Kawakami, and Gammill labs for helpful discussions in group meetings, and Sara Stahley for comments and suggestions. We thank Danelle Devenport for plasmids and keratinocytes and the University of Minnesota Genomics Center for their assistance and expertise in sequencing. Generative AI was used to improve code and original text (ChatGPT-6, Claude-5.5).

---

**Figure S1.**
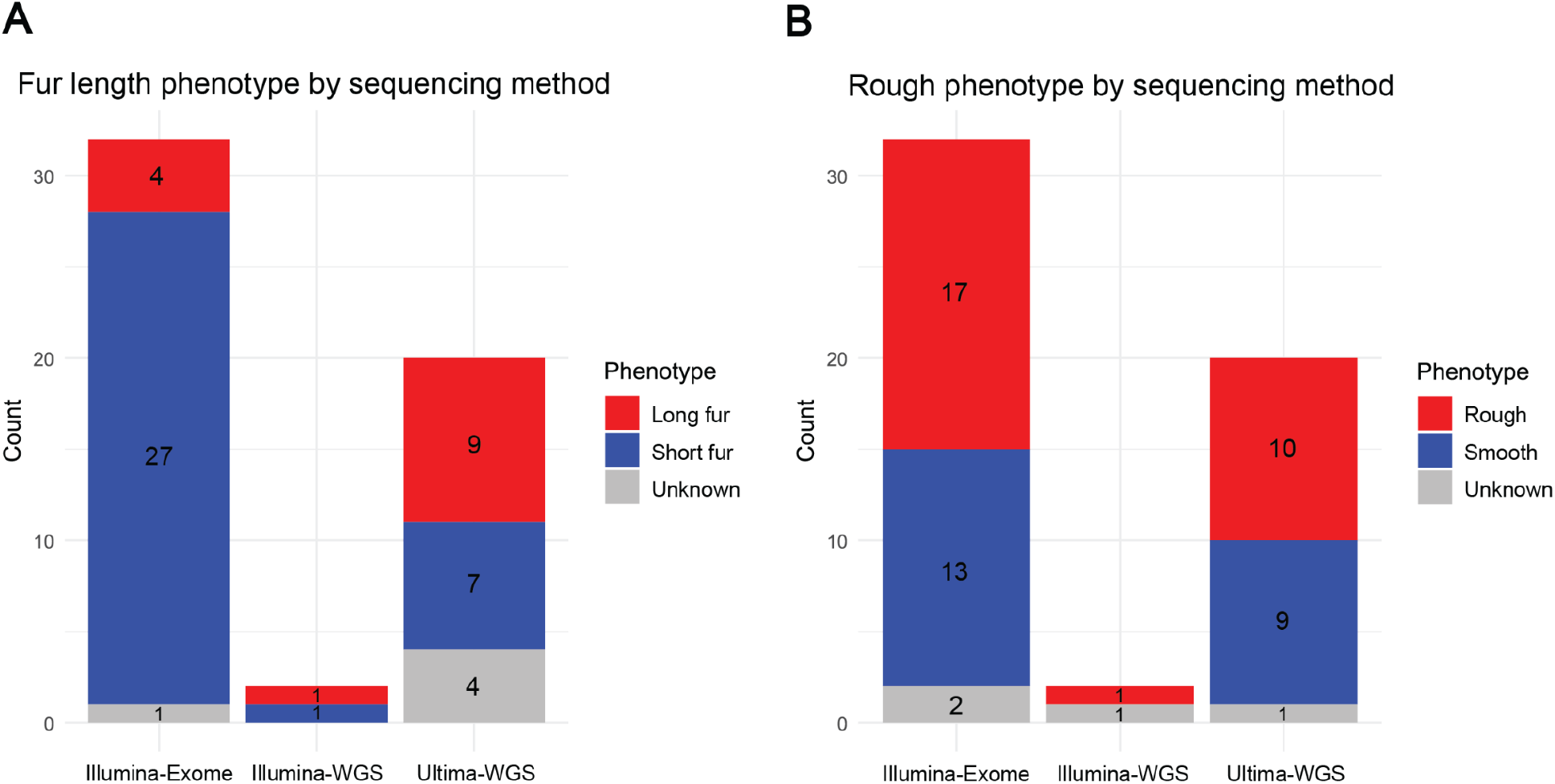
Proportion of sample phenotypes for each sequencing method. (A-B) Stacked bar chart showing sample proportions by sequencing method for long fur (A) and rough coat (B). Blue represents normal phenotypes, red represents the trait of interest, and grey represents unknown phenotypes.

**Figure S2.**
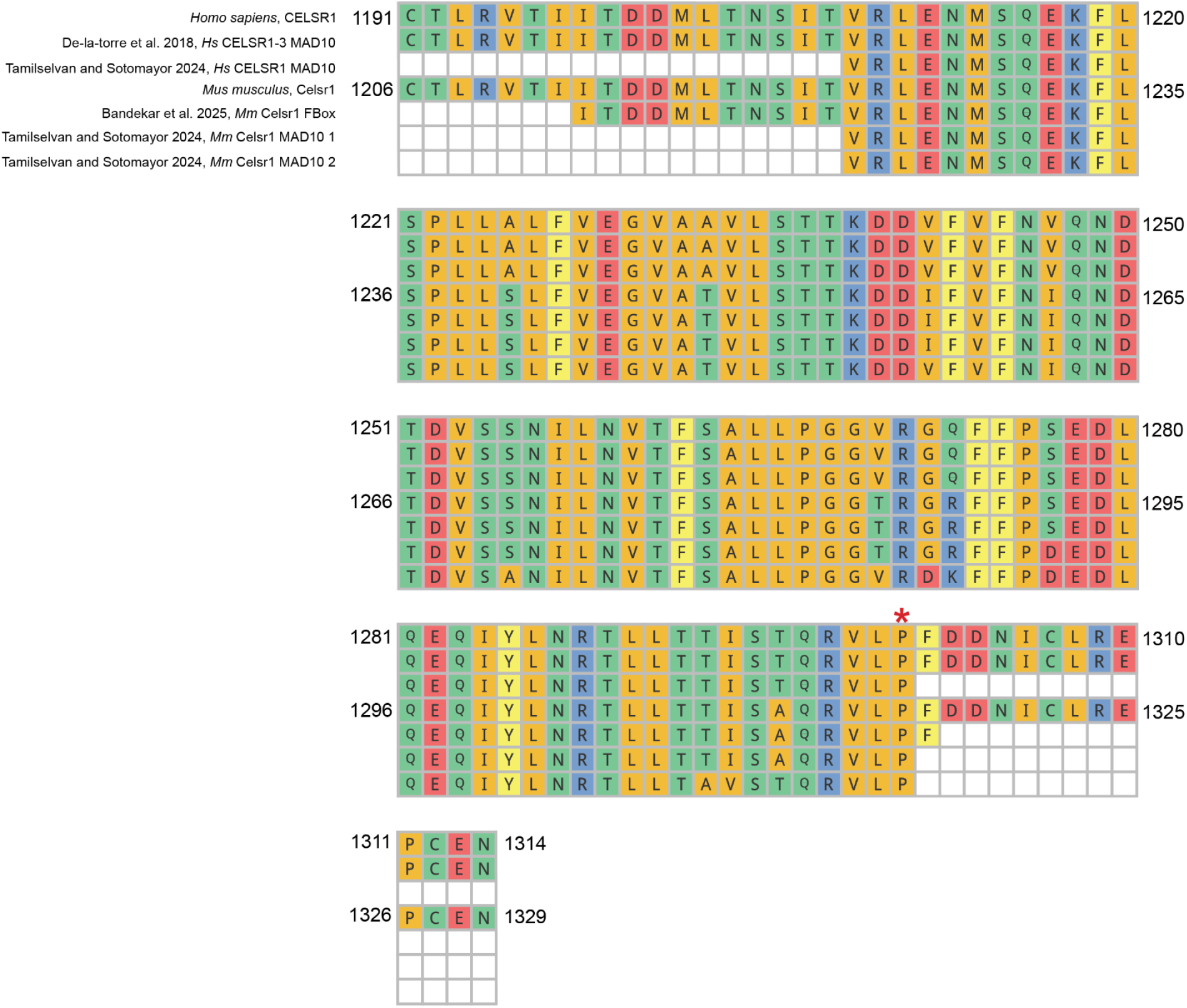
Alignment of Celsr1 MAD10/FBox sequences. Amino acid alignment of human CELSR1 (UniProt Q9NYQ6), human MAD10, mouse Celsr1 (UniProt O35161), mouse FBox, and mouse MAD10. Amino acids are colored by chemical property. Amino acid positions not included in the reference domain sequence are shown as white squares. The proline insertion identified in this study lies within the reported MAD10 and FBox domains and is adjacent to the proline marked by the red asterisk.

**Figure S3.**
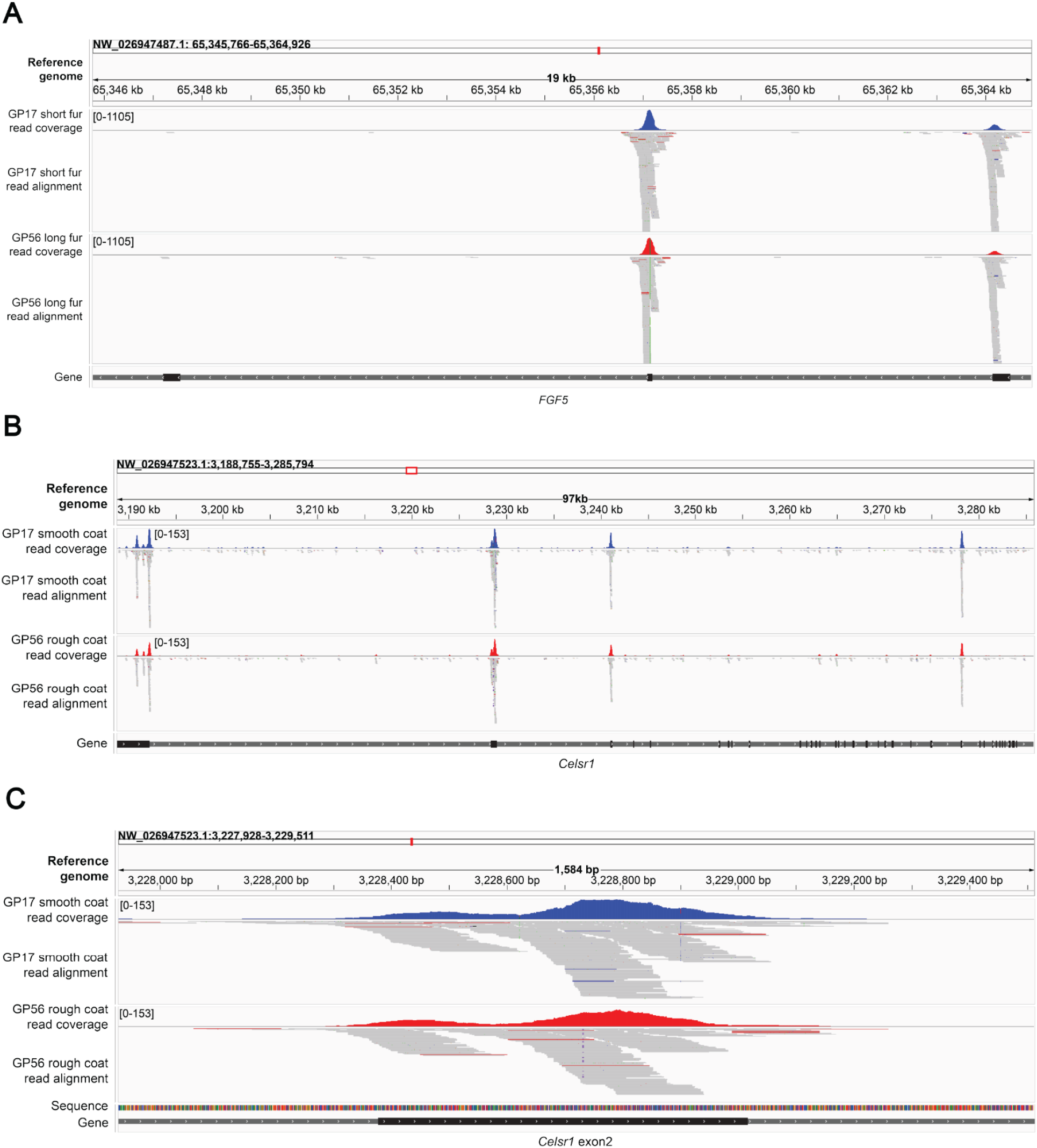
Coverage of *FGF5* and *CELSR1* by human exome capture. Integrative Genomics Viewer (IGV) views of read coverage and alignments for short-fur (blue) and long-fur (red) samples at *FGF5* (A), and smooth-coat (blue) and rough-coat (red) samples at *CELSR1* (B, C). Genomic coordinates and coverage scales are indicated in each panel. Gene exons are shown as black rectangles, with white arrows indicating strand orientation. (A) *FGF5* exons 1 and 2 are well covered, whereas exon 3 is not. (B) Coverage across *CELSR1* is variable, with some exons, including exon 2, well covered and others poorly covered. (C) Enlarged view of *CELSR1* exon 2.

